# Growth-state as a determinant of *in vivo* virulence of *Mycoplasma pneumoniae*

**DOI:** 10.64898/2026.10.06.757197

**Authors:** Jeremy M. Miller, Morgan L. Hunte, R. Grace Ozyck, Edan R. Tulman, Patrick L. Pagano, Kyunghyun Lee, Steven M. Szczepanek, Arlind B. Mara, Steven J. Geary

## Abstract

*Mycoplasma pneumoniae* is a leading cause of community acquired pneumonia and bronchitis in people, particularly children and young adults. Like many bacterial pathogens, *M. pneumoniae* can grow in biofilm towers on both biotic and abiotic surfaces. Current models indicate that planktonic *M. pneumoniae* are simply individual cells that are dispersed from biofilm towers due to physical shear forces, and do not represent a *bona fide* planktonic growth state. However recent data indicate that these fractions may be physiologically distinct and may differentially influence disease outcome. We observed that experimental infection of mice with inoculum containing both adherent and non-adherent fractions of *M. pneumoniae* produced more severe lung pathology than inoculum prepared from *M. pneumoniae* adherent biofilms alone. Direct comparison of experimental infection with titer matched non-adherent (P-Mp) and adherent (B-Mp) fractions revealed that P-Mp produces more severe lung pathology compared to infection with B-Mp. Furthermore, experimental infection with a lower dose of P- Mp resulted in similar lung pathology relative to B-Mp infected mice. This increase in pathology was not associated with differences in bacterial burden, bacterial dispersal during inoculum preparation, nor due to differences in the phenotype of host immune responses. Comparative proteomics revealed that certain known and putative virulence factors associated with immune evasion are upregulated in P-Mp, potentially explaining the difference in virulence between the two fractions. Furthermore, comparative proteomics revealed distinct metabolic phenotypes between the two fractions, consistent with distinct planktonic and biofilm growth states, indicating that P-Mp likely represents a *bona fide* planktonic growth state. Collectively, these data implicate the growth state of *M. pneumoniae* as a determinant of *in vivo* virulence for this atypical human pathogen.

## INTRODUCTION

Lower respiratory tract infections, such as community acquired pneumonia, are the fourth leading cause of death in the world and are estimated to be responsible for approximately $15 billion in direct treatment costs^1–3^. *Mycoplasma pneumoniae* infections account for up to 40% of the cases of community acquired pneumonia, and recent studies place its burden at 3-6 million infections (up from a previous estimate of 2 million) and 100,000 adult hospitalizations annually in the United States alone^4–11^. More recently, *M. pneumoniae* was identified as the cause of the mysterious cases of pediatric pneumonia dubbed the ‘white lung syndrome’ that emerged in China, Europe and United States during the winter of 2023, indicating that *M. pneumoniae* infections have been on the rise worldwide^12–16^. *M. pneumoniae* infections are also associated with the exacerbation, and even predisposition to asthma in children. Coinfections with COVID-19 and influenza are now commonly encountered, and they can result in severe outcomes including death^17–25^. Patients hospitalized with *M. pneumoniae* and COVID-19 co-infections have longer length of hospital stay, ventilator dependence, and higher mortality than patients with COVID-19 alone^23,26^. Co-infection with *M. pneumoniae* and other respiratory viruses, such as adenoviruses, are associated with the progression of *M. pneumoniae* infection to a treatment refractory stage, resulting in longer hospital stay, increased steroid use, and increased need for supplemental oxygen therapy^27–29^.

Despite its significant impact on public health, we remain ill-prepared in our fight against *M. pneumoniae* outbreaks. Mycoplasmas are atypical bacteria that lack a peptidoglycan cell wall, which makes them pan resistant to all β-lactam antibiotics such as penicillins, cephalosporins, cephamycins, monobactams, carbapenems and carbacephems^11^. This innate resistance significantly reduces the arsenal of antibiotics we have to combat *M. pneumoniae* infections, and growing resistance to previously effective macrolide antibiotics raises significant concerns over our ability to control this pathogen.

Further exacerbating the issue of antibiotic resistance, *M. pneumoniae* also form biofilms that are associated with increased resistance to antimicrobial treatment, which often leads to chronic, treatment refractory infections that burden the affected individuals and the healthcare system^30–34^.

Biofilms play a significant role in the virulence of many bacterial pathogens. Biofilms can allow for increased resistance to antibiotics and evasion of the host immune response, allowing them to persist and cause disease in the infected host^35^. In general, chronic infections caused by bacterial biofilms affect roughly 17 million Americans and cause over 550,000 deaths annually in the US alone^36–40^.

Bacterial biofilms are complex communities of microorganisms that adhere to surfaces and are encased in self-produced matrices of extracellular polymeric substances that are typically composed of polysaccharides, proteins, lipids, and extracellular DNA (eDNA), which provide structural stability and protection to the bacterial community^35,36^. Biofilm formation begins when free-floating (planktonic) bacteria attach to a surface, transition to cells that are sessile, and excrete extracellular polymeric substances that shield the bacteria from environmental stressors, antibiotics, and host immune responses^35,36,41^. As biofilms age, the dispersal of biofilm cells occurs via various species-dependent mechanisms^42^. *M. pneumoniae* biofilms are somewhat atypical and divergent from classical model bacteria, owing to the fact that physiologically distinct planktonic cells have not been well defined for this organism. Cell division of *M. pneumoniae* is a concerted effort involving adherence via the organism’s attachment organelle, motility, and FtsZ-mediated constriction of the cell membrane^43^. This means that *M. pneumoniae* predominantly grow within adherent biofilm communities and only possess a transient phase consisting predominantly of individual cells, or cell clusters breaking away from biofilm towers (biofilm breakaways, non-adherent cells). These breakaways exist in suspension for a short period of time before re-adhering and establishing the formation of new biofilms^33^. The mechanisms by which non-adherent *M. pneumoniae* cells are generated via biofilm breakaway are poorly understood. The current models indicate that pieces of *M. pneumoniae* biofilms slough off as a result of physical shear stress from fluids, indicating that the sloughed aggregates/non-adherent *M. pneumoniae* maintain the characteristics of biofilms and do not represent differentiated planktonic phases^44^. However, recent work has demonstrated that these fractions of *M. pneumoniae* possess different metabolomic profiles, indicating that the two growth states may be biologically distinct^45^. Recent *in vitro* investigations modeling *M. pneumoniae* biofilm infections in submerged BEAS-2B bronchial epithelial tissue culture revealed that biofilm breakaway aggregates are important for establishing infection in eukaryotic cells^34^. Due to recent developments, we investigated if these breakaways truly represent passively sloughed off pieces of *M. pneumoniae* biofilms, or whether they are actively derived, *bona fide* planktonic bacteria that are phenotypically distinct from those found in the biofilms. Additionally, we compared the virulence and disease outcomes between these two fractions hereafter referred to as P-Mp (non-adherent fraction) and B-Mp (adherent fraction).

## RESULTS

### Infection of mice with non-adherent *M. pneumoniae* results in more severe lung pathology than infection with the biofilm counterpart

We had observed from our historical animal experiment data that mice infected with *M. pneumoniae* inoculum prepared from cultures with mixed adherent (hereafter referred to as B-Mp) and non-adherent (hereafter referred to as P-Mp) growth states developed more severe pulmonary lung lesions than mice infected with only *B-Mp* at 4 days post-infection(**Supplementary Figure S1**). Previous work by other groups demonstrated that increased dispersal of *M. pneumoniae* inoculum prior to infection resulted in better take of infection and increased persistence of *M. pneumoniae* in the lungs of experimentally challenged mice^46^. Collectively considered, these data made us curious about the role that different growth states of the pathogen could play in the *in vivo* virulence of *M. pneumoniae.* To directly address this question, we prepared titer matched inoculum from solely P-Mp, mixed P-Mp and B- Mp (M-Mp), and solely B-Mp. We then infected mice through intranasal instillation, performed necropsies 4 days post-infection, assessed lung pathology, and quantified bacterial burdens in the lung. We found that mice infected solely with P-Mp had more severe lung pathology when compared to mice infected solely with B-Mp, while mice infected with M-Mp had intermediate pathology (**Figure 1A, D**). No significant differences were observed in lung bacterial burdens based on viable bacteria recovered from the lungs of infected animals (**Figure 1B**), and lesion scores poorly correlated with bacterial burden (**Figure 1C**). Altogether, these data indicated that P-Mp infection produced more severe lung pathology than B-Mp in a manner that appeared to be unrelated to lung bacterial burden.

**Figure 1.**
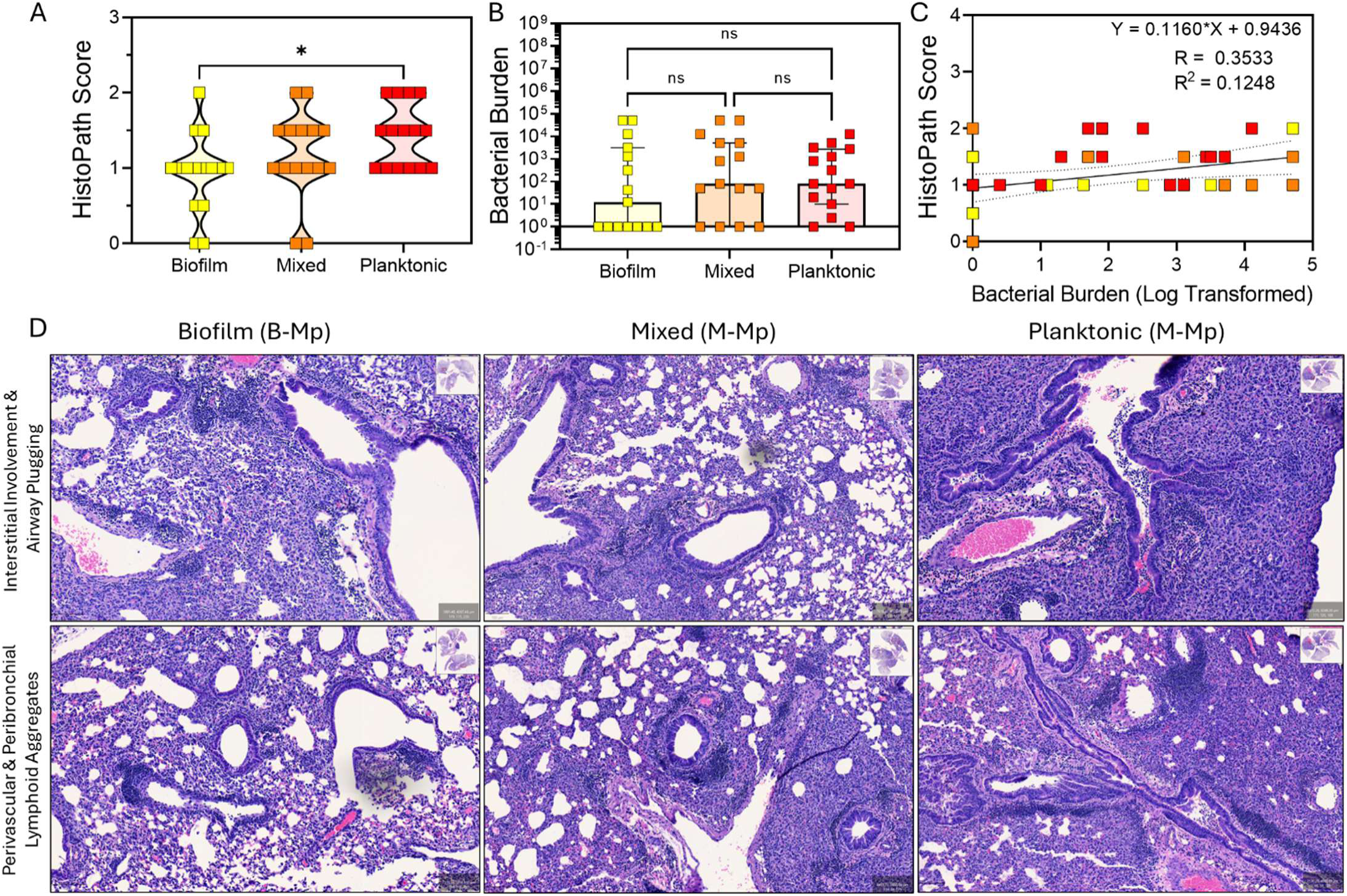
P-Mp infection produces more severe disease than B-Mp infection. (**A**) Histopathological lung lesion scores and (**B**) bacterial burdens of infected animals. (**C**) Linear regression analysis displaying relationship between histopathological score and bacterial burden and (**D**) Representative histopathology from infected animals.

### Increased dispersal of inoculum does not account for heightened virulence of non-adherent *M. pneumoniae*

Previous studies indicated that increased dispersion of bacteria during inoculum preparation improves persistence of *M. pneumoniae* in the lungs of infected mice, indicating that our observed increase in P-Mp virulence may be due to increased dispersal of bacterial cells within this fraction^46^. To assess this hypothesis, we prepared biomass-matched (measured via OD620) inoculum containing solely P-Mp or inoculum containing solely B-Mp. We then sonicated a portion of both fractions to obtain biomass-matched dispersed fractions (P-Mp (S) and (B-Mp (S)). Sonication resulted in an 8.5- to 10-fold increase in viable bacterial counts over non-sonicated fractions (**P-Mp**: 7.44x10^6^ CCU/50uL versus **P- Mp(S)**: 7.44x10^7^ CCU/50uL| **B-Mp:** 1.22x10^8^ CCU/50uL versus **B-Mp(S)**: 1.04x10^9^ CCU/50uL) as measured using the most probable number (MPN) method using serial dilutions and assessment of growth via color changing units (CCUs). Despite this increase in CCUs, dispersal of inoculum through sonication did not impact the ability of the different fractions to cause disease, as all fractions produced lung lesions that were not statistically different from each other 4 days post-infection (**Figure 2A,C, D F**). This indicated that the increased virulence of P-Mp was not due to increased dispersal of bacterial cells in this fraction. We also observed no differences in bacterial burdens in the lungs or bronchoalveolar lavage fluid (BALF) of mice infected with the sonicated versus non-sonicated fractions (**Figure 2B, E**).

**Figure 2.**
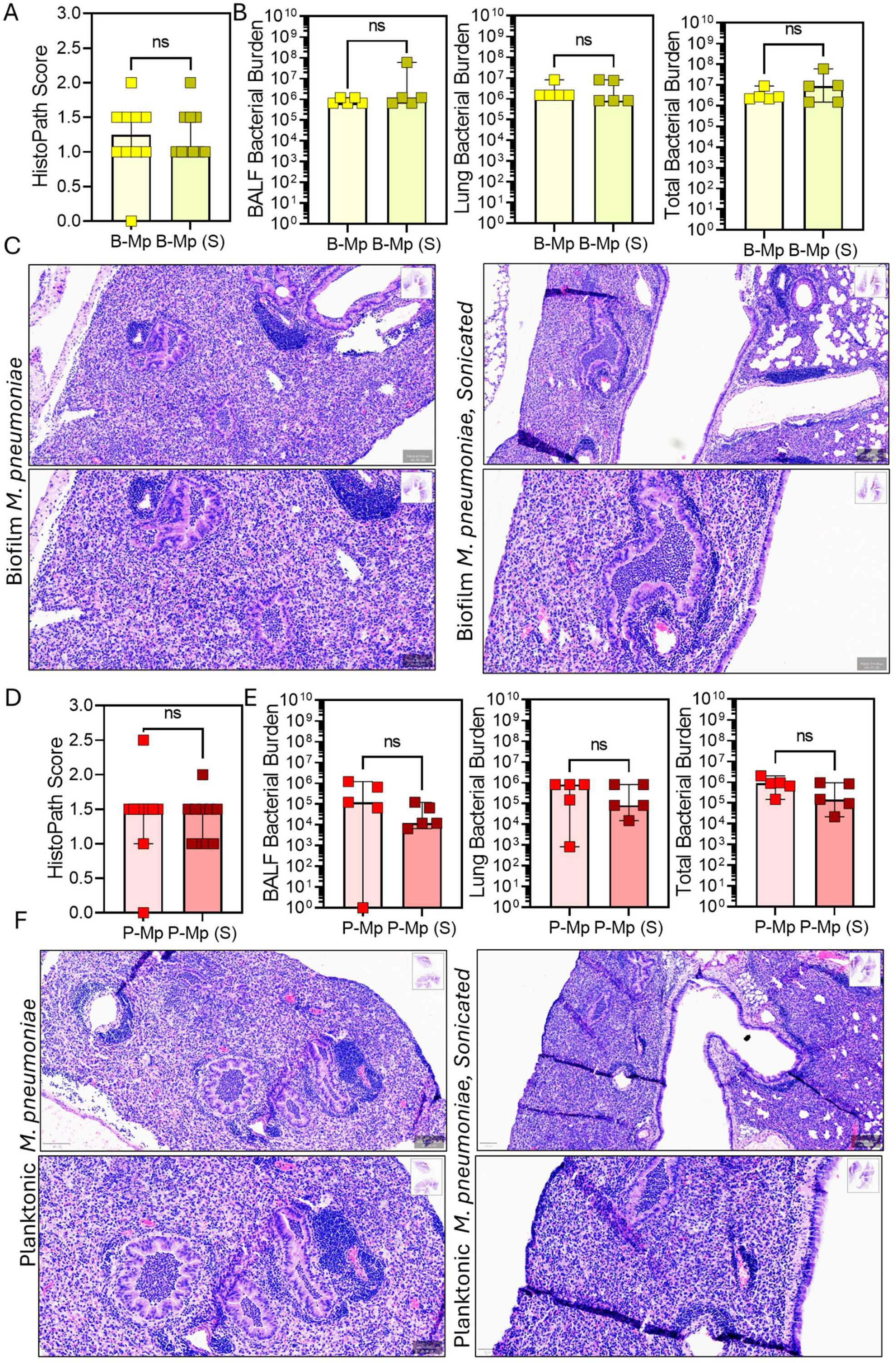
Elevated virulence of P-Mp is not due to increased dispersal of bacterial units. (**A**) Histopathological lung lesion scores and (**B**) bacterial burdens of infected animals in bronchoalveolar lavage fluid (BALF), lung tissue, or combined total. (**C**) Representative histopathology from animals infected with B-Mp fractions. (D) Histopathological lung lesion scores and (E) bacterial burdens of infected animals in bronchoalveolar lavage fluid (BALF), lung tissue, or combined total. (F) Representative histopathology from animals infected with P-Mp fractions.

Mice infected with inoculum prepared from P-Mp developed lung pathology that was similar in severity to that observed in mice infected with B-Mp, despite bacterial titters of the inocula being 13- to 16-fold lower in the P-Mp fractions than in the B-Mp fractions. This further underscored the elevated virulence of P-Mp, which does not appear to be linked to increased dispersal of inoculum (Supplementary **Figure S2**).

### Infection with non-adherent *M. pneumoniae* induces host immune responses that are elevated in magnitude, but are generally similar in phenotype to those induced by infection with adherent *M. pneumoniae*

Having established that increased dispersal of inoculum did not explain the increased virulence of P-Mp, we wished to assess if the difference in virulence could be explained by differential host responses to the different fractions. To this end, we performed a longitudinal study where we tracked the host responses to infection through extensive immunophenotyping of recruited leukocytes collected via bronchoalveolar lavage over a 21-day time course. Every 3 days post infection, we collected bronchoalveolar lavage fluid (BALF) and immunophenotyped airway inflammation using our previously developed full-spectrum flow cytometry panel that can identify with high accuracy the following cell populations: *leukocytes, erythrocytes (RBCs)*, *non-leukocytes/epithelial*, *neutrophils*, *eosinophils*, *tissue-resident alveolar macrophages (ResAMs)*, *monocyte-derived macrophages* (*moMacs*) including monocyte-derived *alveolar macrophages* (*moAMs*), *natural killer cells* (NK), *CD4^+^ T cells*, *CD8^+^ T cells*, *γδ T cells*, *NKT-like cells*, and *B cells* (**Supplementary Figure S3**)^47^. Following collection and analysis of all timepoints via flow-cytometry, the data were integrated, dimensionality reduction was performed using Uniform Manifold Approximation and Projection (UMAP), and both cell frequencies and counts were assessed using conventional gating strategies^48^. Analysis of UMAP-generated leukocyte clusters revealed that airway inflammation matured similarly over time in both P-Mp and B-Mp infected animals, though P- Mp infected animals appeared to have a higher number of leukocytes in the airways across several timepoints, though these did not always reach statistical significance (**Figure 3** and **4**, **Supplementary Figures S4** and **S5**). The most striking difference was the significant elevation of moMacs, which were increased in both frequency and count in *P-Mp* at day 6 post-infection (**Figure 3** and **4**, **Supplementary Figures S4** and **S5**). Other leukocyte populations also differed in count at various timepoints (**Figure 4**, **Supplementary Figure S5**), but they did not differ in frequency (**Supplementary Figure S4**), indicating that host responses induced by P-Mp and B-Mp infections are very similar in phenotype, though generally higher in magnitude in mice infected with P-Mp, which is consistent with the observed increase in lung pathology. It is unclear whether the significant differences in frequency and magnitude of moMacs at day 6 post-infection are important for the observed increase in virulence of P-Mp. While moMacs are typically more pro-fibrotic and can contribute to lung pathology through degradation of extracellular matrix and replacement of functional lung tissue with fibrotic scar tissue, previous reports also indicate that macrophages may play a role in the clearance of *M. pneumoniae* from the lungs of infected mice^49,50^.

**Figure 3.**
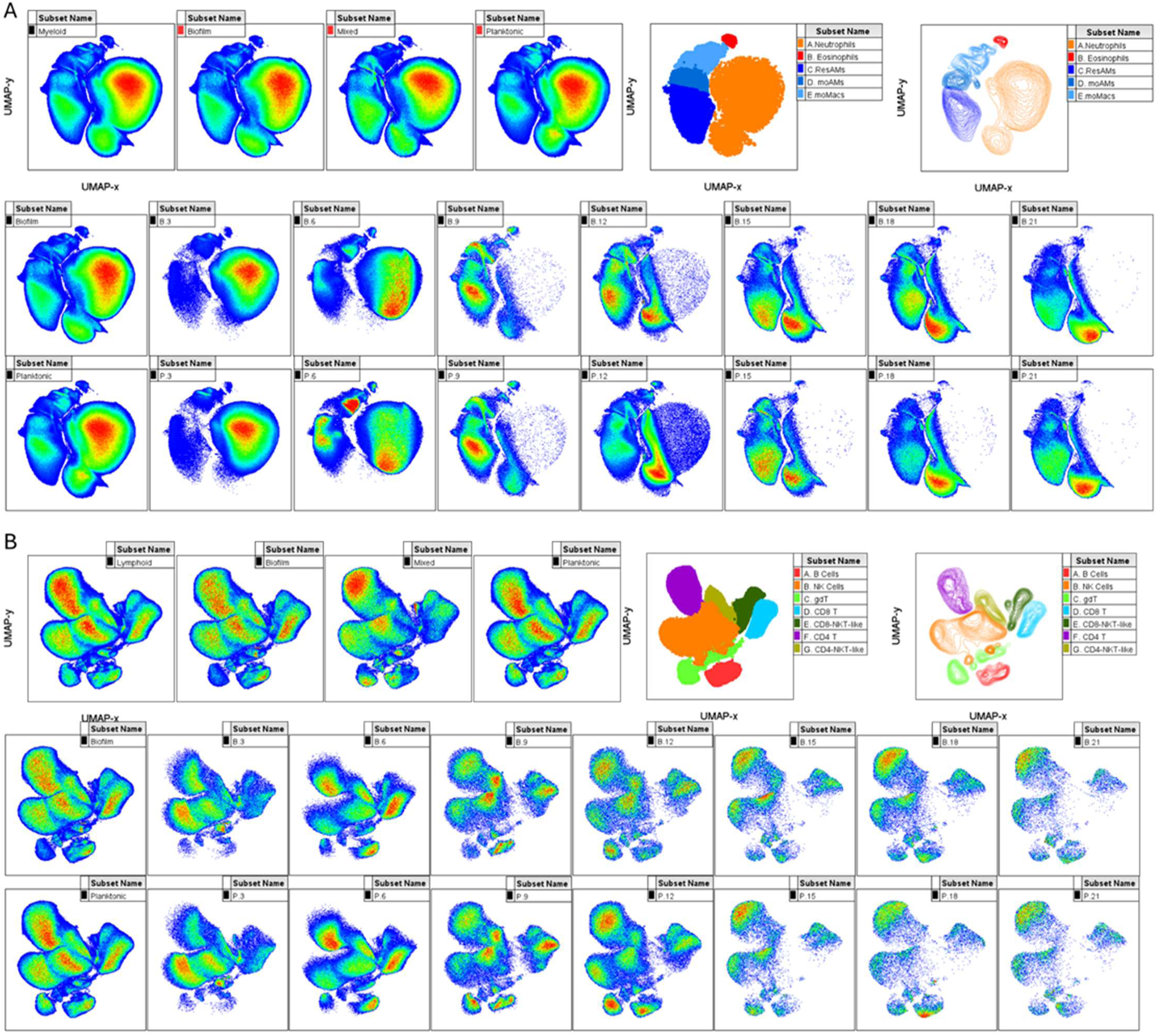
Elevated virulence of P-Mp is unlikely to be due to differential host responses to infection by the different fractions. Longitudinal immunophenotype of host immune responses (**A**: Myeloid Cells, **B**: Lymphoid Cells) in the airways of infected mice displayed as UMAP-dimensionality reduced pseudo color density plots.

**Figure 4.**
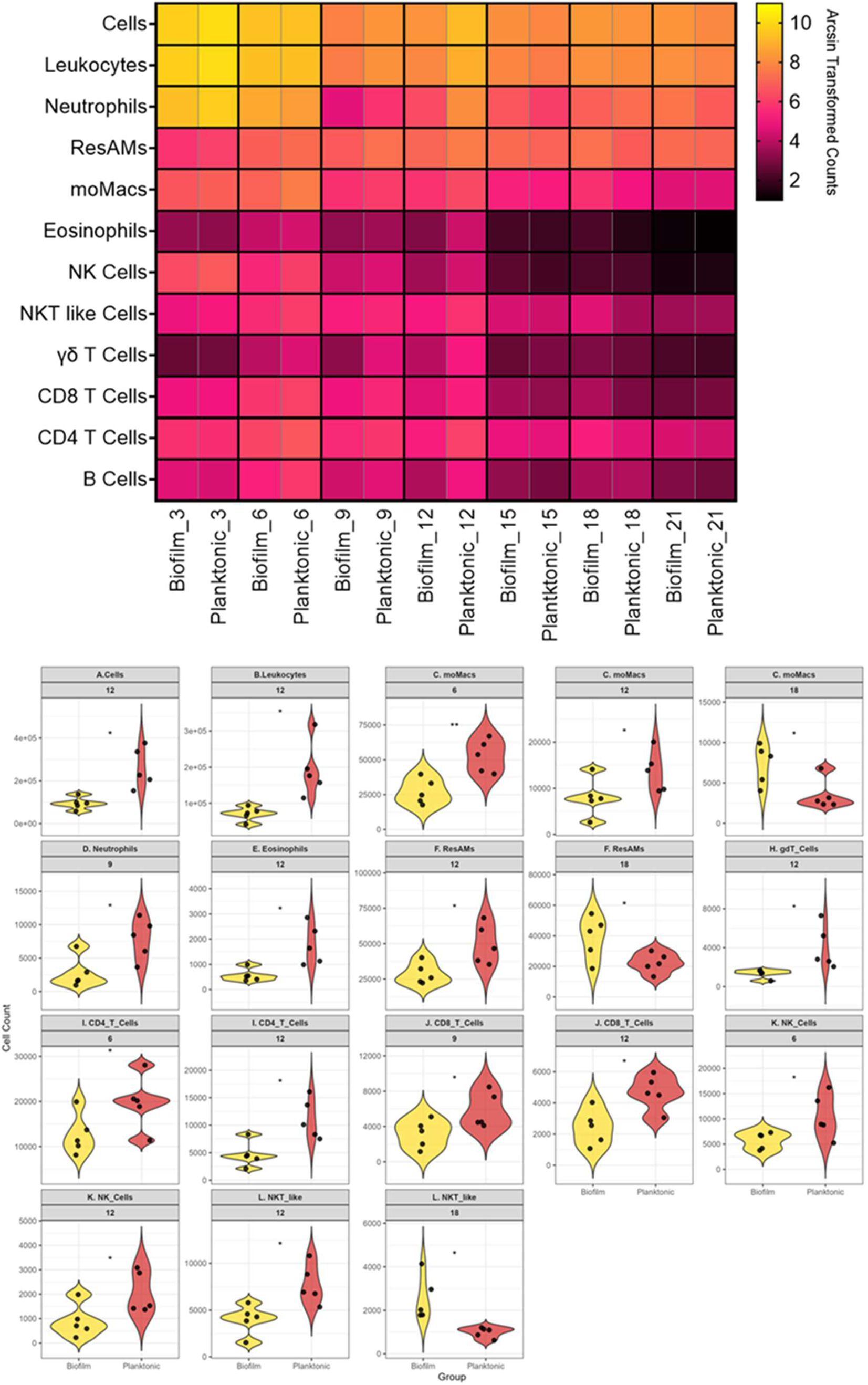
Leukocyte composition of airway inflammation changes similarly over the course of infection in both P-Mp and B-Mp infected mice. (A) Heatmap indicating relative makeup and (B) significantly different absolute counts of leukocytes in mice infected with B-Mp versus P-Mp.

While this study did not distinguish whether this protective role was played by tissue-resident macrophages, or those derived from inflammatory monocytes, current knowledge prevents us from making any definitive statements regarding the meaning of the elevated moMacs in P-Mp infected mice. Bacterial burdens in the BALF and lung tissue of infected animals did not differ over time between mice infected with P-Mp or B-Mp, indicating that the fractions do not appear to differ in their ability to colonize the lung (**Figure 5**). It is, however, worth noting that *M. pneumoniae* was more consistently recovered from P-Mp infected mice at 15DPI (100% of animals versus 20% of animals positive), which hints at the possibility that P-Mp may be better poised to colonize the lungs. Furthermore, the increase in moMacs at day 6 post-infection in the P-Mp infected animals does not appear to correlate with reduced bacterial burden, which indirectly indicates that this population is likely not contributing to bacterial clearance, but rather it is associated with increased lung pathology. However, given that the majority of the differences in host responses are in magnitude (elevated counts) rather than phenotype (frequencies), it is our interpretation that the observed increase in the virulence of P-Mp over that of B-Mp is not due to the induction of differential host responses. More detailed mechanistic studies are needed to clarify the role that moMacs may be playing in this phenomenon.

**Figure 5.**
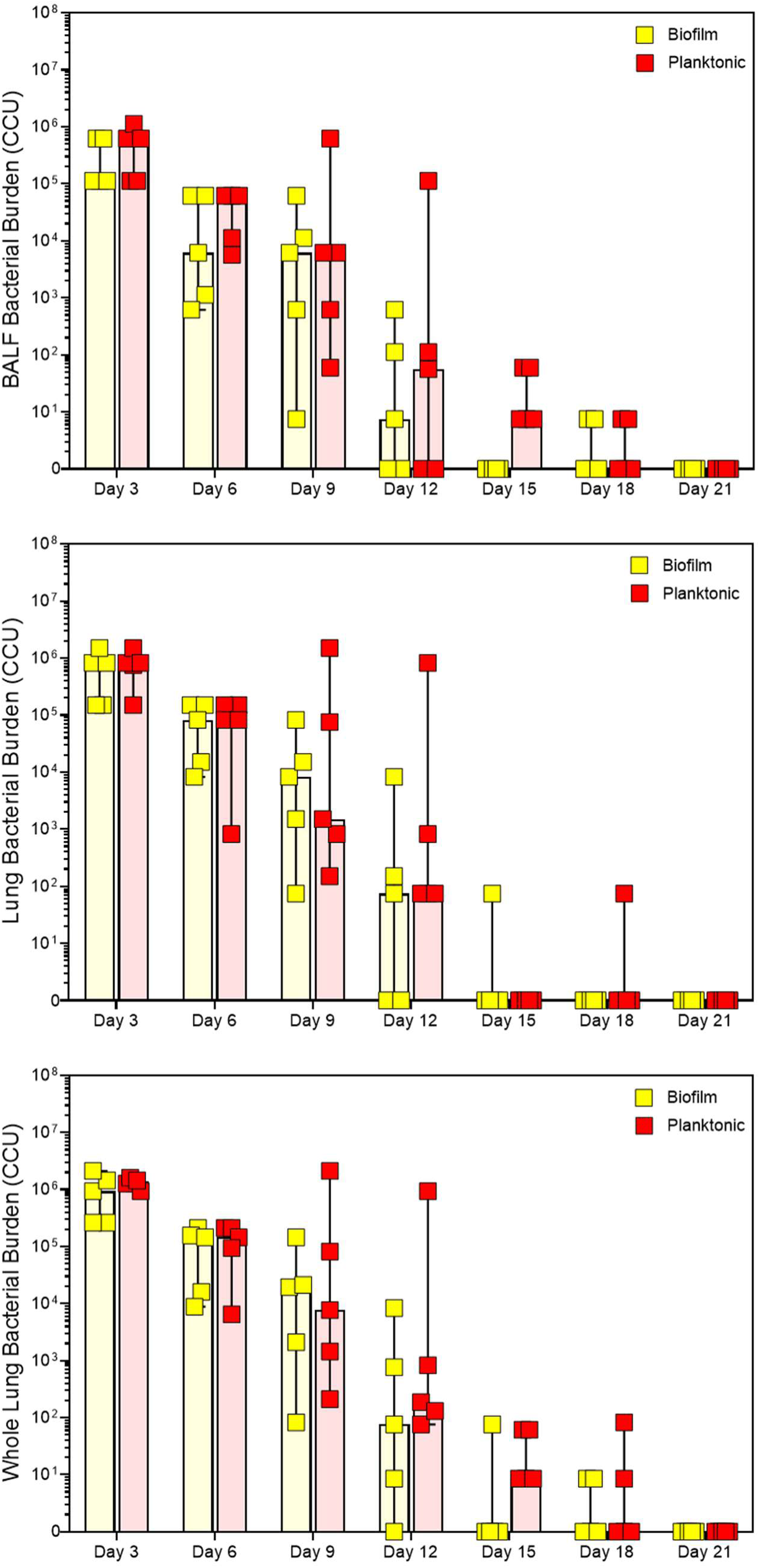
Longitudinal recovery of viable *Mycoplasma pneumoniae* from lungs of mice infected with P-Mp versus B-Mp. Color changing units of viable Mp in (**A**) BALF, (**B**) Lung Tissue, and (**C**) combined total recovered from lungs of infected animals.

### Comparative proteomic profiling reveals divergent metabolic, virulence, and stress response programs in different *M. pneumoniae* growth states

Having shown that increased inoculum dispersal and differential host responses are unlikely to explain the difference in the virulence of P-Mp and B-Mp, we performed label-free proteomics via LC/MS- MS, to identify phenotypic differences between these growth stages. Differential abundance analysis identified 128 proteins that were significantly differentially expressed, with 41 proteins being significantly enriched in P-Mp, and 87 being enriched in B-Mp (**Figure 6**, **Supplementary Table S1**). Four proteins were found to have very high fold-change differences between the fractions. Of these four, enriched in the B- Mp were the MgpC family cytadherence protein (ortholog of MPN102; log2FC of −29.90), DUF1600 domain containing protein (MPN274; log2FC of −24.99) and Holliday junction resolvase RuvX (MPN418; log2FC of -9.97). Enriched in P-Mp was phosphopantetheine-binding protein (MPN406; log2FC of 30.66). An in-depth description of proteomic differences between the B-Mp and P-Mp states is discussed in the supplementary file. Briefly, proteins involved in cell division, energy metabolism, translational machinery, immune evasion, cytadherence and virulence factors, the glycerol-to-hydrogen peroxide virulence axis, and oxidative stress defense were among those differentially enriched in each fraction. Proteomic profiling indicated that P-Mp and B-Mp likely represent disparate phenotypes, consistent with what is reported between planktonic and biofilm growth states of other bacterial species (see detailed supplementary discussion).

**Figure 6.**
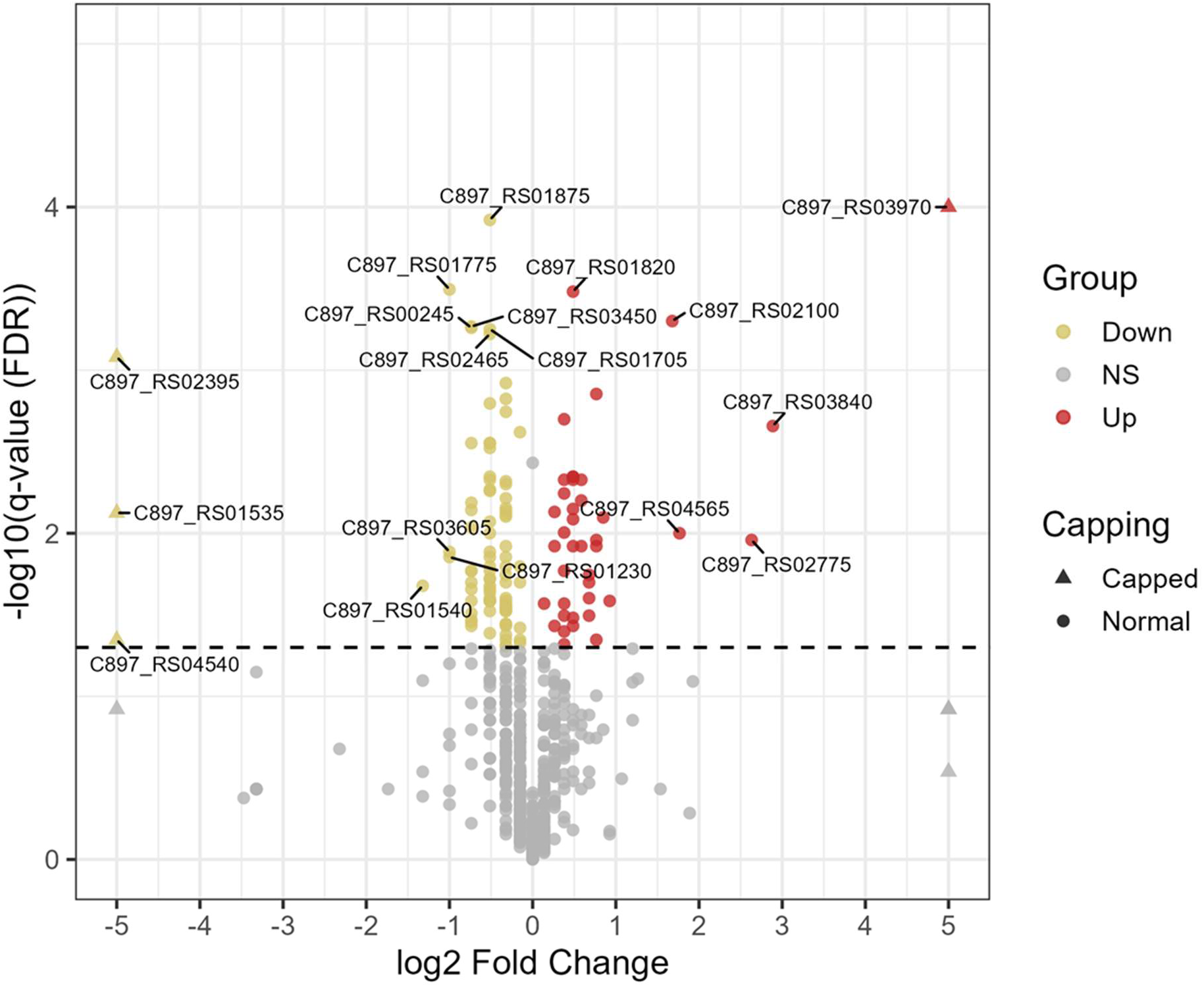
Proteomic profiling reveals divergent phenotype between P-Mp and B-Mp growth states. Volcano plot indicating differentially enriched proteins in the P-Mp fraction (red) and B-Mp fraction (yellow). Triangles indicate proteins that were present in one fraction and absent in the other, with capped Log2 fold change values to improve visualization.

More relevant for the virulence axis discussed in this manuscript, several known and putative virulence factors were upregulated in P-Mp, which can potentially explain the heightened *in vivo* virulence of this fraction. These include the thermonuclease family protein (MPN133), the membrane nuclease (MPN491), the IgG-blocking protein M IbpM (MPN400), oligoendopeptidase F (MPN197), dihydrolipoyl dehydrogenase (MPN390, DLD, DLDH, lipoamide dehydrogenase, *lpd*, E3*, pdhD*), nucleotide exchange factor GrpE (MPN120), and P200 (MPN567). MPN133 is a cytotoxic nuclease that can be internalized into cells and cause cell death and pathology and MPN491 is a secreted nuclease that mediates immune evasion by digesting neutrophil extracellular traps^108,109^. IbpM mediates antibody escape by sequestering IgM, IgG and IgA antibodies, and a mutant in this gene has displayed reduced cytotoxic capabilities^110^.

MPN197 is a surface accessible peptidase that may be involved in *M. pneumoniae* surface protein presentation and has been found to serve as a negative regulator of biofilms^111,112^. A homolog of this protein in a related Mycoplasma species was found to cleave bradykinin, substance P, neuropeptide Y, and neurokinin A, potentially implicating it as a virulence factor as well^113^. MPN390 is a critical enzyme involved in the pyruvate dehydrogenase complex and has been demonstrated to be a key virulence determinant for multiple organisms, including the closely related *Mycoplasma gallisepticum*. It has also been identified as a plasminogen-binding protein and is thus also implicated in adherence^114–117^. GrpE is a co-chaperone with DnaK and has been found to be located on the surface of other Mycoplasma species and contribute to adhesion by interacting with plasminogen and other extracellular matrix components^118^. Further demonstrating the increased ability of P-Mp to adhere to host components, is the upregulation of the adherence associated protein P200 (MPN567) which is required for attachment to differentiated bronchial epithelium and for the gliding motility of the organism^119^. The enriched P200 in P-Mp relative to B-Mp is consistent with other literature on *M. pneumoniae* biofilms indicating that P200 is downregulated over time as biofilms mature^44^.

Also noteworthy was the significant upregulation of the *phosphopantetheine binding protein* (AcpP, MPN406) in P-Mp, which was almost entirely absent in the B-Mp. MPN406 is an acyl carrier protein that is a component of fatty acid synthesis (FAS), non-ribosomal polypeptide synthesis (NRPS), and polyketide synthesis(PKS)^132^. Acyl carrier proteins are implicated in bacterial virulence through 4’- phosphopantetheinyl transferases (PPTases), which are enzymes that post-translationally attach a phosphopantetheine moiety (Ppant) from CoA onto carrier proteins. This attachment activates these carrier proteins for biosynthesis of virulence-related molecules in other bacterial pathogens such as *Mycobacteria* spp. *Salmonella enterica*, pathogenic *E. coli*, *Burkholderia*, *Acinetobacter*, *Pseudomonas aeruginosa,* and even some fungi^133–146^. *M. pneumoniae* lacks the ability to synthesize fatty acids *de novo,* and instead extracts them from the host cells through the actions the lipid scavenger protein P116 (MPN_213)^147^. However, *M .pneumoniae* still possesses the acyl carrier protein synthase (AcpS |MPN_298) which is the 4’-phosphopantetheinyl transferase (PPTase), and the acyl carrier protein AcpP (MPN406) which provides phosphopantetheine as a cofactor for delivery acyl groups to phospho- or glycolipids, which these genes may be involved in the synthesis of^133,146^. Both genes are suggested to be essential, and data are limited on whether they play a role in the virulence of *M. pneumoniae*. *M. pneumoniae* also lacks the genes required for the synthesis of traditional siderophores, and it is not currently known whether the production of the CARDS toxin or other *M. pneumoniae* virulence factors relies on AcpP activity. However, many Mycoplasma proteins are known to moonlight as virulence factors and given that the AcpS/AcpP axis is associated with *in vivo* virulence of other bacterial pathogens, it is tempting to speculate that AcpP may contribute to, either directly or indirectly, to the heightened virulence of P-Mp.

Strikingly, the *CARDS toxin*, an ADP-ribosylating and vacuolating exotoxin with established roles in airway inflammation and pathology, the hydrogen sulfide producing *cysteine desulfurase HapE*, the lipid acquisition protein P116, nor the P1 and P30 adhesins were differentially expressed between P-Mp and B- Mp, suggesting that the expression of these major virulence factors was not regulated by the growth state of *M. pneumoniae*^43,147–150^. It is worth noting, however, that different P1 adhesin loci, including the main gene (MPN141) as well as the repetitive elements, were differentially detected across growth states but showed no consistent trends across replicates, indicating the P1 adhesin dynamics in the different life stages of *M. pneumoniae* are more nuanced than can be resolved with simple proteomic analysis.

## DISCUSSION

Historically, the field has treated planktonic *M. pneumoniae* as a transient byproduct of biofilm shear rather than a physiologically and phenotypically distinct growth state. However, our data align with emerging metabolomic evidence that *M. pneumoniae* possesses a planktonic state that is phenotypically distinct from its biofilm counterpart. Notably, our data demonstrate that *Mycoplasma pneumoniae* possesses a phenotypically distinct planktonic growth state (P-Mp) that differs significantly in its capacity to cause disease *in vivo* when compared to the biofilm fraction. Planktonic *M. pneumoniae* was found to produce more severe disease than titer matched biofilm counterpart (**Figure 1**). Biofilms dispersed through sonication failed to recapitulate this elevation in virulence, and lower doses of non-adherent *M. pneumoniae*, which we argue to be *bona fide* planktonic *M. pneumoniae*, produced similar disease to higher dose biofilm *M.pneumoniae* regardless of inoculum dispersal(**Figure 2**). This finding challenges the current assumption that planktonic cells in *M. pneumoniae* cultures merely represent passively sloughed off biofilm fragments and instead supports the existence of a *bona fide* planktonic growth state with unique biological properties. Although only a transient growth state, our comparative proteomics analysis indicates that planktonic *M. pneumoniae* represents a divergent phenotype from that of *M. pneumoniae* biofilms.

Many have tried to compare planktonic and biofilm fractions of bacteria, but the methods that have been used thus far almost never allow for both fractions to have been grown in the same vessel, at the same time. Thus, our method for preparing P-Mp and B-Mp uniquely allowed us to directly compare between the two growth states while limiting confounding variables. Comparative proteomics of the two different fractions delineate the fractions as phenotypically distinct. Most notably, the B-Mp were enriched for proteins involved in adherence, membrane flux, glycolysis, fatty acid/membrane synthesis, stress responses, and biofilm formation. In contrast, the P-Mp were upregulated for proteins involved in virulence, cell division, energy generation, and phenotype switching(**Figure 6, Table 1, Supplementary Table S1**).

The increase in MgpC family cytadherence protein in B-Mp could be a mechanism by which the cells within the biofilm attach to one another as adhesins are commonly associated with biofilm formation^151^. We observed upregulations in proteins involved in glycolysis, which is common under oxygen depletion, upregulation in proteins involved in hydrogen peroxide production, and a multitude of stress responses in the biofilm fraction. Oxygen depletion, presence of hydrogen peroxide, and starvation have also all been reported as signals for biofilm dispersal^152^. Specifically, it is plausible that B-Mp are responding to stress induced by their own hydrogen peroxide production, thus triggering an environment that sets the stage for planktonic cell release.

We see several pieces of evidence that biofilm dispersal in *M. pneumoniae* is an active process.

The Holliday junction resolvase RuvX cleaves branched DNA, which is interesting because eDNA of biofilms is often organized into structures resembling Holliday junctions^91,153^. *M. pneumoniae* biofilms are not currently believed to contain eDNA, as DNase I treatment was not able to disrupt *M. pneumoniae* biofilms as it does with other bacterial biofilms^33,34^. However DNaseI cannot act on Z-DNA, which eDNA of bacterial biofilms is frequently composed of^33,154,155^. Indeed, it has been shown that biofilms of *Helicobacter pylori* contain eDNA that is not susceptible to DNaseI, and biofilms of a related *Mycoplasma* species (*M. hyopneumoniae*) contain eDNA that can be disrupted by benzonase, which is a DNase that can act on Z-DNA^94,156,157^. Thus, *M. pneumoniae* biofilms may contain eDNA made of Z-DNA, and the B-Mp may be using RuvX to modify the biofilm matrix as it grows. Interestingly, the MPN491 nuclease upregulated in the planktonic fraction is able to cleave Z-DNA, which is what composes DNA in neutrophil extracellular traps (NETs)^158^. It is possible, therefore, that MPN491 may be one of the endogenous nucleases that cleaves *M. pneumoniae* biofilm eDNA in order to mediate the dispersal of planktonic cells from biofilm towers. Such a mechanisms has been shown for other bacterial species that form biofilms such as *Staphylococcus* species^159^. Therefore, it is possible that MPN491, which was enriched in the P-Mp, mediates escape from biofilm towers while simultaneously allowing for immune escape through NET degradation. The cytotoxic thermonuclease MPN133, also enriched in the P-Mp, could also contribute to eDNA breakdown and biofilm dispersal while also serving as a virulence factor mediating host damage. The enrichment of oligoendopeptidase F in the P-Mp may also be playing a role in the break-away of planktonic cells from *M. pneumoniae* biofilms through digestion of surface proteins holding the biofilm together, as has shown to be possible through proteinase K treatment being able to disrupt *M. pneumoniae* biofilms^33^. It should be noted that oligoendopeptidase F has also been noted to serve as a negative regulator of bacterial biofilms, which further supports this hypothesis^112^. Interestingly, oligoendopeptidase F may additionally function as a virulence factor as a homolog of it in a related *Mycoplasma* species can bind bradykinin, substance P, neuropeptide Y, and neurokinin A, which implicates the factor in immune modulation^113^.

Several putative and known virulence factors were significantly upregulated in P-Mp, including the immunoglobulin blocking IbpM, and several proinflammatory lipoproteins. These factors drive immune evasion, and potentially immunopathology, that could explain the enhanced virulence of P-Mp. One particularly intriguing finding is the strong upregulation of the phosphopantetheine-binding protein AcpP (MPN406) in the P-Mp. Mutation of AcpP in *P. aeruginosa* was found to increase biofilm formation, suggesting that MPN406 serves as a negative regulator of biofilm formation^160^. As noted in the results, AcpP is also associated with various virulence associated mechanisms in other bacteria. Although a direct link between AcpP and *M. pneumoniae* virulence is not yet understood as it is in other bacteria, the near-exclusive expression of AcpP in P-Mp raises the possibility that phosphopantetheinylation- dependent processes contribute to virulence in this growth state. While speculative, this represents a promising avenue for future mechanistic studies.

It is important to note that the increased virulence of the planktonic growth state is likely multifactorial. Given that we ruled out improved dispersal, improved persistence, and major differences in host immune responses as mechanisms for this increase in virulence, we propose that the answer lies within the proteomic differences. Based on literature and our data presented here, we propose a putative model which could explain the distinct virulence phenotypes of *M. pneumoniae* growth-states which we illustrate in **Figure 7**. Our work builds on a model proposed by the Balish lab which proposed that biofilm breakaways (which we here demonstrate are likely true planktonic *M. pneumoniae*) are more cytotoxic and produce more virulence factors while biofilms are more poised for persistence and display resistance to clearance by antibiotics and complement^33,34^. Our results identified several virulence-associated proteins enriched within the planktonic fraction, many of which may be additionally involved in biofilm dispersal. It is likely that these proteomic differences partially explain the observed increase in virulence, but because virulence is multifactorial, it remains unclear how much individually identified factors contribute to the observed phenotypes. It remains unknown whether the degree of upregulation of these factors is high enough to cause the increase in virulence observed, and it is also likely that many of the uncharacterized proteins that are differentially enriched contribute to this phenotype.

**Figure 7.**
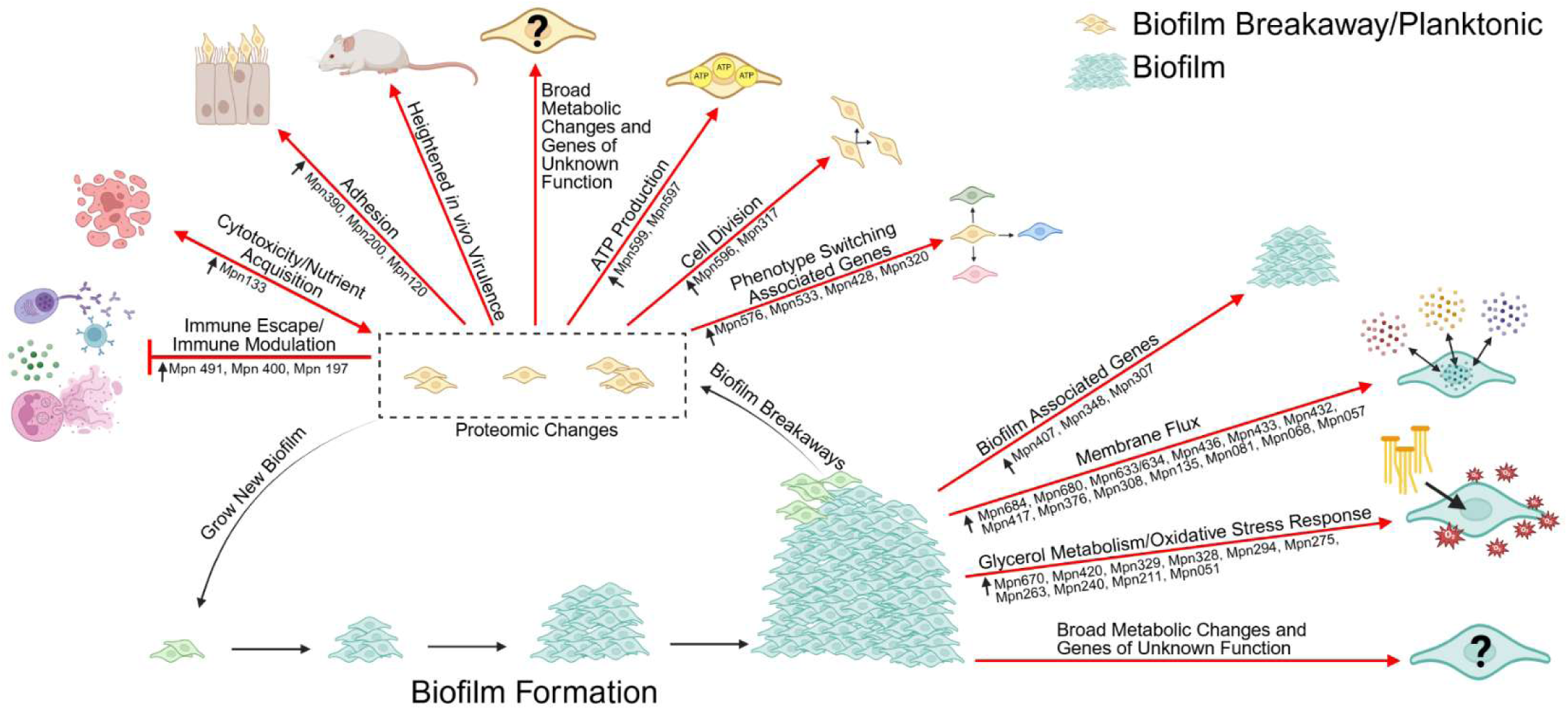
Illustration of the proposed expanded model illustrating the potential reasons for the observed enhanced virulence of the P-Mp fraction.

## CONCLUSION

Our data provide the first *in vivo* evidence that planktonic *M. pneumoniae* represent a distinct, highly virulent physiological state. We show that the enhanced virulence of planktonic *M. pneumoniae* is not attributable to bacterial load, inoculum dispersal, or major differences in host responses, but instead correlates with a proteomic profile enriched for metabolic activity, immune evasion, and cytotoxic potential. These findings fundamentally reshape our understanding of *M. pneumoniae* biology and highlight growth-state heterogeneity as a key driver of virulence for this insidious human pathogen.

## METHODS

### Bacteria and Inoculum Preparation

Low passage *Mycoplasma pneumoniae* strain PI1428 was used for all experiments reported herein. The fractions used for animal experiments were generated at the same timepoint using the same initial seed stock for maximal within-experiment consistency. *M. pneumoniae* were inoculated into T-175 flasks with Fortified Commercial (FC) medium supplemented with 20% heat-inactivated horse serum, and cultured at 37°C until a shift in pH was observed, indicating growth. The supernatant was discarded to remove debris and potentially dead *M. pneumoniae,* and fresh FC growth medium was added. Flasks were again incubated at 37°C until a shift in pH was observed, indicating growth. *M. pneumoniae* were then harvested using a cell scraper, and the collection was divided among new T-175 flasks filled with fresh FC growth medium. Flasks were incubated at 37°C until a pH shift was observed. Supernatant was removed to discard debris and potentially dead cells, and fresh FC medium was added. Once a pH shift was then observed, harvest differed by intended bacterial fraction. If used to generate mixed inoculum, both the adherent and non-adherent fractions were collected, centrifuged (all centrifugation steps were performed at 10,000 xg for 10 minutes at 4°C), combined, and resuspended in fresh FC media. OD620 measurements were taken to estimate CFU, samples were centrifuged, resuspended at 5x10^8^ CFU/mL in FC medium, and tubes were frozen at −80°C for future use. For generation of planktonic (non-adherent) and biofilm (adherent) inocula, the supernatant was removed as planktonic cells, centrifuged, and frozen at −80°C as pellets. Fresh FC media was added to the flasks for another round of growth. This step is required to generate sufficient inoculum as the planktonic phase is a small percentage (∼10%) of the *M. pneumoniae* biomass. Once a pH shift was observed, supernatant was removed as planktonic cells, combined with previously frozen pellets, centrifuged, and resuspended in FC medium. OD620 measurements were taken to estimate CFU, and samples were then centrifuged, resuspended at 5x10^8^ CFU/mL in FC medium, and tubes were frozen at −80°C for future use. Biofilm cells were scraped off the same flasks the planktonic *M. pneumoniae* were taken from. They were scraped into fresh FC medium, centrifuged, combined, and resuspended. OD620 measurements were taken to estimate CFU, samples were then centrifuged, resuspended at 5x10^8^ CFU/mL in FC medium, and tubes were frozen at −80°C for future use.

### Inoculum preparation

Sufficient tubes needed to infect the designated number of mice were thawed, placed into fresh FC medium, and incubated at 37°C with shaking at 180 rpm for five hours. Tubes were then centrifuged and resuspended in fresh FC medium. OD620 measurements were taken to estimate CFU, and samples were then centrifuged and resuspended in fresh FC medium at a concentration of 2x10^9^ CFU/mL (1x10^8^ CFU/50uL) for infections. For studies where sonicated fractions were generated, these inocula were divided in half, with one set being sonicated for one second at 20% amplitude using a (**insert machine specific info here**) probe sonicator to generate sonicated, dispersed fractions. Serial 10-fold dilutions were performed on a 96-well plate to quantify the functional titer of bacteria using the Most Probable Number (MPN) method.

### Animal Experiments

7-week-old JAX® BALB/cJ (000651) female mice (The Jackson Laboratory, Farmington CT) were acclimated for one week prior to experimentation. Food and water were provided *ad libitum* across the entire study period. Work was conducted in accordance with our animal protocols approved by the University of Connecticut Institutional Animal Care and Use Committee. For infections, mice were anesthetized using vaporized isoflurane and were intranasally inoculated with 50 uL of FC medium containing a total of 1x10^8^ CFU of M. pneumoniae. Following infections, inocula were serially diluted to confirm equivalent viable titer between groups using the MPN method. Mice were necropsied at four days-post-infection for all studies except the timecourse study, where mice were necropsied every three days-post-infection out to 21 days-post-infection.

Outputs for the first study (biofilm vs mixed vs planktonic) were histopathology via lung lesion scores and lung snip bacterial burdens. Outputs for the second study (sonicated vs non-sonicated) were lung lesion scores, whole lung bacterial burdens, and bronchioalveolar lavage fluid (BALF) bacterial burdens.

Outputs for the third study (timecourse) were whole lung bacterial burdens, BALF bacterial burdens, and flow cytometric analysis of BALF.

### Histopathology

For histopathology, lungs were allowed to fix in 10% neutral buffered formalin for 24 hours before being placed into cassettes and submitted to the Connecticut Veterinary Medical Diagnostic Laboratory for embedding, sectioning, and staining with hematoxylin and eosin. Slides were evaluated as previously described^161,162^. In brief, lesions were scored as follows: 0 (no visible lesions), 1 (mild lesions), 2 (moderate lesions), 3 (marked lesions), 4 (severe lesions), with half scores (e.g. 1.5) given to samples between two scores.

### Bacterial Burdens

For lung snip bacterial burdens, small snippets of the right caudal lobe were placed into FC medium on ice. Tubes were then vortexed for two minutes and incubated at 37°C for three hours. Medium from the tube was then filtered through a 0.45 µM Whatman™ Uniflow™ disposable sterile syringe filter (Cytiva, Marlborough MA). Four-fold serial dilutions were performed into FC medium with samples run in duplicate to establish color changing units (CCUs). Values reported were an average of duplicate wells.

For whole lung bacterial burdens, lungs were removed and placed into 3 mL of FC medium on ice. Lungs were removed from tubes and chopped with scissors until a chewing-gum-like consistency was produced. Lung bits were then put back into the same tube and vortexed for 30 seconds. Samples were incubated at 37°C for one hour. Samples were then put through a 70 µM cell strainer (Corning Life Sciences, Tewksbury MA) to remove lung pieces. Samples were then put through a 0.45 µM Whatman™ Uniflow™ disposable sterile syringe filter (Cytiva, Marlborough MA). Ten-fold serial dilutions were performed into FC media with samples run in duplicate to establish color changing units (CCUs). Values reported were an average of duplicate wells.

For BALF bacterial recovery, the BALF sample was centrifuged at 450 xg, 4°C, for five minutes and 200 uL of supernatant from each BALF sample was added to 3 mL of FC medium. Samples were then filtered through a 0.45 µM Whatman™ Uniflow™ disposable sterile syringe filter (Cytiva, Marlborough MA). Ten-fold serial dilutions were performed into FC medium with samples run in duplicate to establish color changing units (CCUs). Value reported was an average of duplicate wells. Reported, “total recovery” is the result of summating an individual animal’s whole lung burden and BAL burden.

### Bronchioalveolar Lavages

Bronchioalveolar lavages were performed by inserting an PrecisionGlide™ 18 G, 1” needle (Becton, Dickinson and Company, Franklin Lakes NJ) attached to a 3 mL syringe (Becton, Dickinson and Company, Franklin Lakes NJ) filled with 1.5 mL of sterile phosphate buffered saline (PBS) into the trachea of the mouse. Slowly, 1 mL of the PBS was dispensed into the trachea. This fluid was then pulled back out. This was repeated two more times using the same liquid for three total washes. The BALF was then placed into a tube on ice.

### Flow Cytometry

For flow cytometric analysis of BALF, we utilized our established protocol. Dimensionality reduction was performed via UMAP, and all analyses were conducted on FlowJo version 10.10.0.^47^

### Proteomics

Mycoplasma pneumoniae PI1428 was inoculated into FC medium inside a T175 flask (Thermo Fisher Scientific, Waltham MA). This grew at 37°C until media began acidifying (as determined by a color change from red to orange). Supernatant was discarded. Fresh FC medium was added and the flask was incubated at 37°C until media began acidifying. The flask was then scraped and divided amongst 10, T175 flasks. FC medium was added to each flask and incubated at 37°C until media began acidifying.

Supernatant was removed and pooled from flasks 1-3, 4-7, and 8-10 as Planktonic samples 1, 2, and 3 respectively. Planktonic samples were centrifuged, supernatant was discarded, and samples were resuspended in PBS. PBS was added to flasks and flasks were scraped using a sterile 39 cm handle, 30 mm Lifter Blade cell scraper (CELLTREAT, Ayer MA). Flasks 1-3, 4-7, and 8-10 were then combined as “Biofilm” samples 1, 2, and 3 respectively. Planktonic and Biofilm samples were re-centrifuged and resuspended in PBS until 3 total washes occurred. Samples were resuspended in 1 mL PBS with 5% SDS. Protein concentrations were determined using Pierce™ BCA Protein Assay Kit - Reducing Agent Compatible (Thermo Fisher Scientific, Waltham MA).

Samples were then processed by using S-Trap columns, where the samples are reduced to break any disulfide bonds, alkylated to modify cysteine sidechains and prevent disulfides from re-forming, and then loaded onto the column and incubated with protease overnight to digest intact proteins into peptides. Peptide products are then eluted off the column and desalted to remove any buffer components or intact protein that may have co-eluted. The samples were quantified by absorbance measurements at 280 nm and a 500 ng aliquot was injected for each sample. We tested a 2-hr gradient based on the expected complexity and the instrument scan speed was sufficient to capture the available sample data over a 2-hr elution. Based on this, the remaining samples were also run at 2 hrs. The data were searched against the annotated *Mycoplasma pneumoniae* PI1428 genome (NZ_CP010538.1). A common contaminants database was also included to account for proteins that can be introduced during sample generation or prep, which includes bovine serum proteins (from cell culture media), keratins from several species (to account for skin and clothing), and common proteases (added during mass spec sample prep). All the results are filtered to a 1% false discovery rate threshold, meaning that out of the total list of proteins, approximately 1% of them are false positive identifications.

### Data Analyses and Statistics

Scaffold version 5 was used to visualize and analyze proteomics data. Relative protein abundances were reflected as Normalized Average Precursor Intensities, and multiple T-tests with the Benjamini-Hochberg false discovery rate correction were run to determine significant differences in proteins expression between the Biofilm and Planktonic fractions. Volcano plots were generated in R version 4.5.2. FlowJo Version 10.1.10 was used to analyze flow cytometry data. GraphPad Prism version

9.0.0 was utilized for statistical analyses of histopathological lesion scores and bacterial burden data. Non-parametric Mann-Whitney U tests were utilized to compare data between two groups, while one-way ANOVA on Ranks (Kruskal-Walis) was used to compare differences in more than two experimental groups given a single independent variable. Time course data were analyzed via a Two-Way ANOVA mixed model. Dunnet and Sidak post-hoc tests for multiple comparisons were run following the appropriate analysis of variance test.

## Author Contributions

Conceptualization (J.M.M, S.M.S, A.B.M., E.R.T., S.J.G.)

Methodology (J.M.M, A.B.M, M.L.H, R.G.O., E.R.T., S.M.S., S.J.G.)

Software C Formal Analysis (J.M.M, A.B.M.) Validation (J.M.M, A.B.M.,)

Investigation (J.M.M, A.B.M, M.L.H, R.G.O.) Resources (S.J.G.)

Data Curation (J.M.M, M.L.H, A.B.M,) Writing – Original Draft (A.B.M)

Writing – Review C Editing (J.M.M, M.L.H, R.G.O, A.B.M, E.R.T, S.M.S, S.J.G.)

Visualization (A.B.M.) Supervision (A.BM, S.J.G.)

Project Administration (A.B.M., E.R.T., S.M.S., S.J.G.) Funding Acquisition (S.J.G.)

---

**Sup. Fig. 1.**
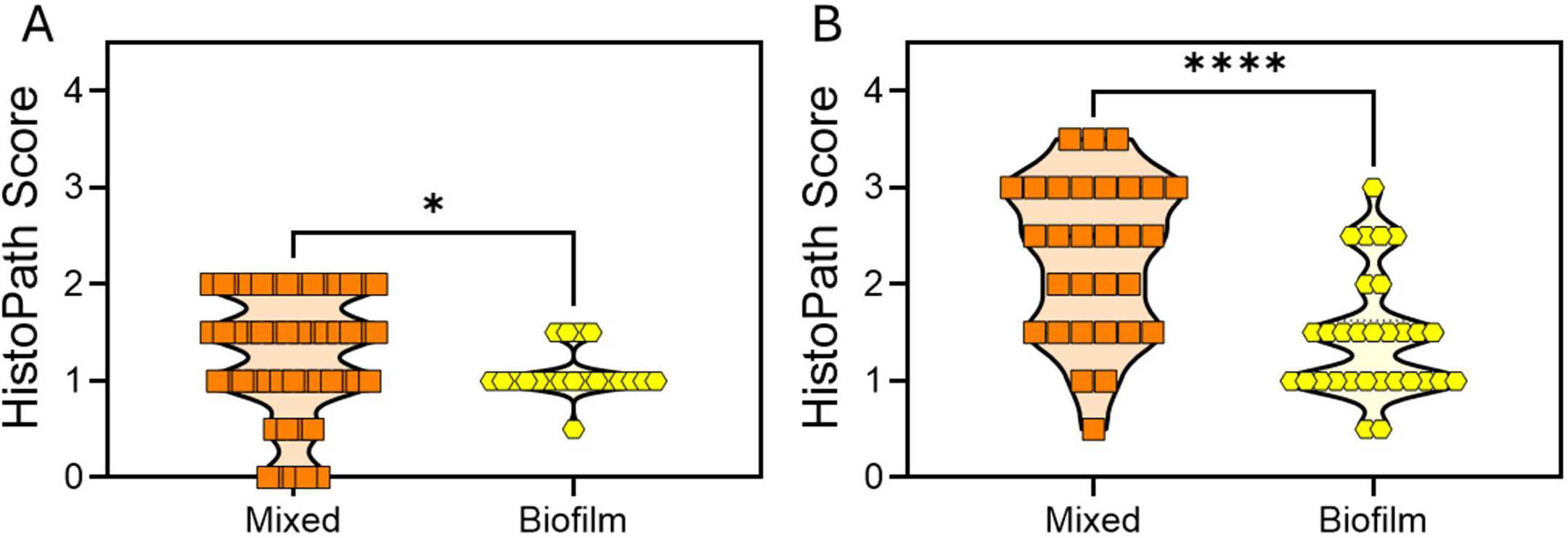
Lab historical data indicating differences in lung pathology induced by infection of mice with mixed *M. pneumoniae* inoculum containing both adherent and non-adherent fractions (biofilm + planktonic), versus fractions containing only adherent/biofilm. (**A**) *M. pneumoniae* infection of naïve mice and (**B**) mice previously vaccinated with *M. pneumoniae* lipid associated membrane proteins experiencing vaccine enhanced disease.

**Sup. Fig. 2.**
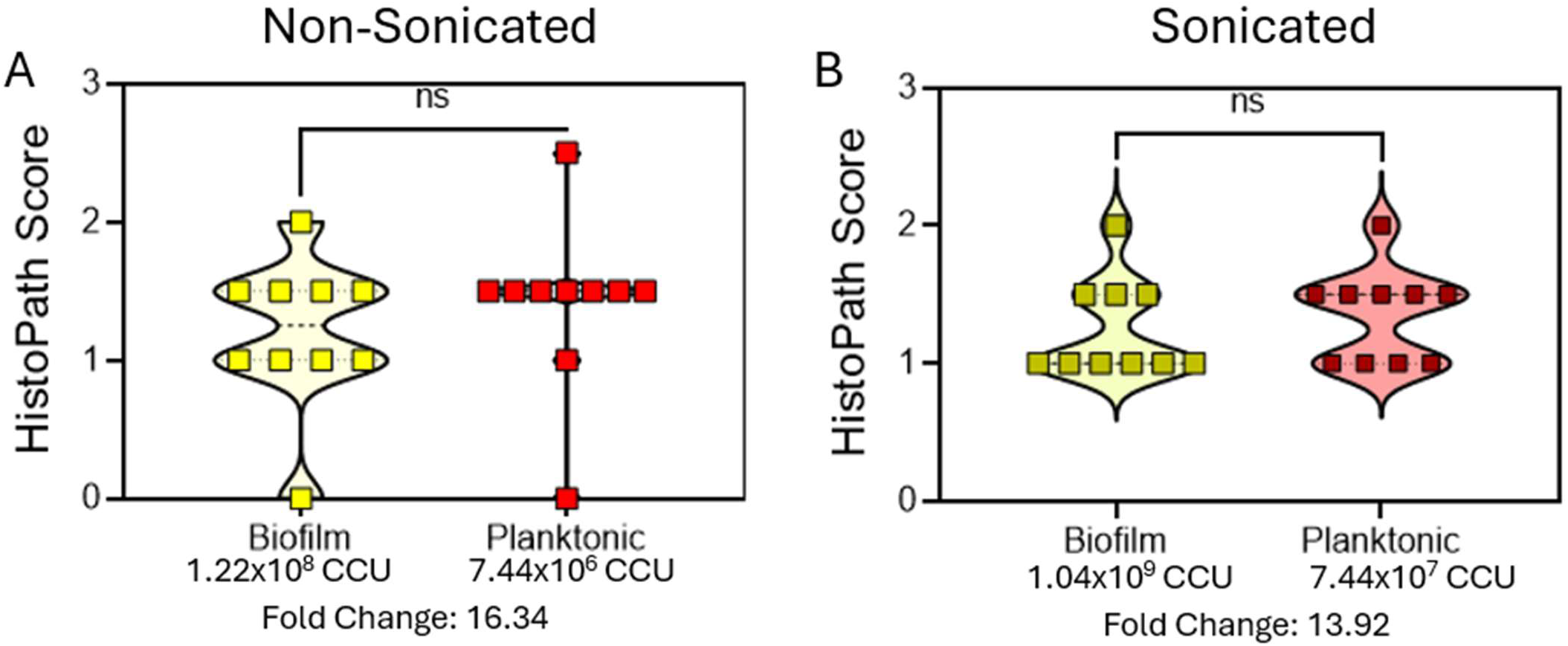
Infection with a lower titer of P-Mp produces similar lung pathology to higher titer B-Mp infection, regardless of dispersal status.

**Sup. Fig. 3.**
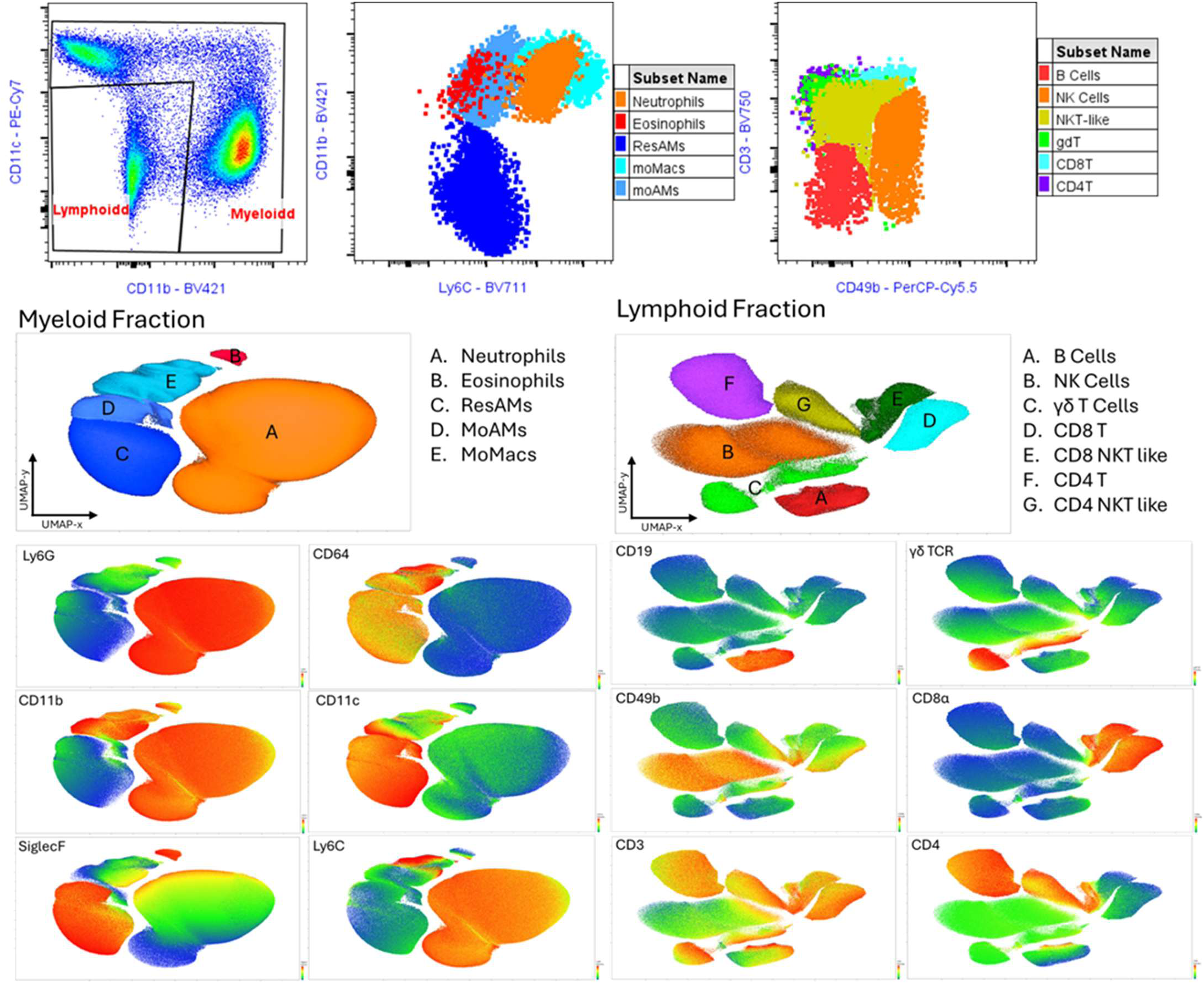
Leukocyte marker validation for UMAP cell cluster identification. **Myeloid Cells:** Neutrophils: CD45^+^Ly6G^+^Ly6C^+^CD11b^+^CD64^-^CD11c^-^ SiglecF^var^, Eosinophils: CD45^+^Ly6G^-^ Ly6C^-^CD11b^+^CD64^-^CD11c^-^ SiglecF^hi^, Resident Alveolar Macrophages (ResAMs): CD45^+^Ly6G^var^Ly6C^-^CD11b^-^CD64^+^CD11c^+^SiglecF^hi^, Monocyte-derived Alveolar Macrophages (moAMs): CD45^+^Ly6G^lo^Ly6C^+^CD11b^lo^CD64^+^CD11c^+^SiglecF^+^, Monocyte-derived Macrophages (moMacs): CD45^+^Ly6G^lo^Ly6C^hi^CD11b^hi^CD64^hi^CD11c^lo^SiglecF^lo^. **Lymphoid Cells (**All on CD45+CD11c-CD11b-CD64-Ly6G- pregate): B Cells: CD19^+^CD3^-^ CD49b^-^ CD4^-^ CD8^-^ γδTCR^-^, NK Cells: CD19^-^CD3^-^ CD49b^+^ CD4^var^ CD8^var^ γδTCR^-^, NKT-like Cells: CD19^-^CD3^+^ CD49b^+^ CD4^var^ CD8^var^ γδTCR^-^, γδ T Cells: CD19^-^CD3^+^ CD49b^-^ CD4^var^ CD8^var^ γδTCR^+^, CD4 T Cells: CD19^-^CD3^+^ CD49b^-^ CD4^+^ CD8^-^ γδTCR^-^, CD8 T Cells: CD19^-^CD3^+^ CD49b^-^ CD4^+^ CD8^+^ γδTCR^-^.

**Sup. Fig. 4.**
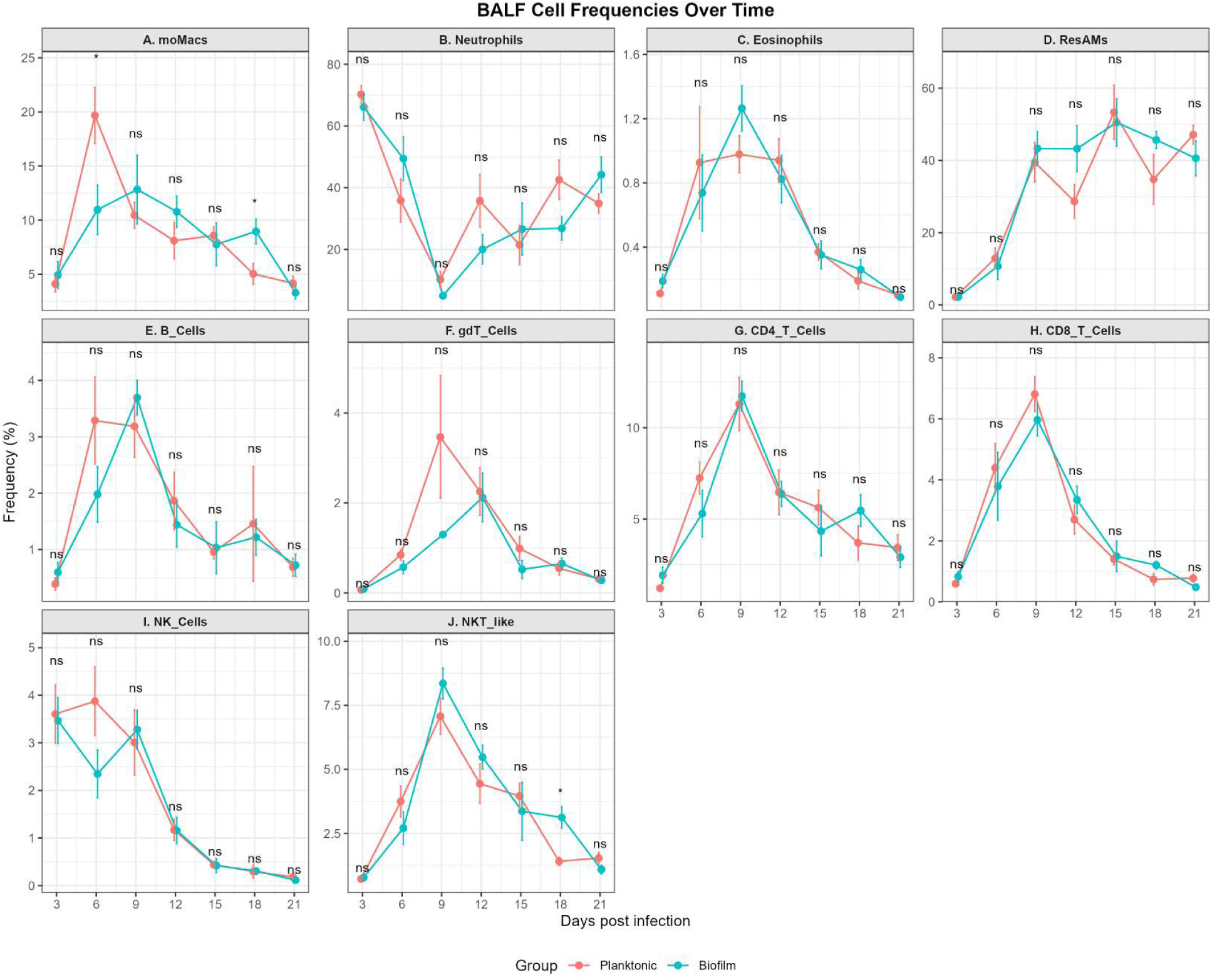
Kinetics of leukocyte frequencies in bronchoalveolar lavage fluid of *M. pneumoniae* infected mice per day post infection.

**Sup. Fig. 5.**
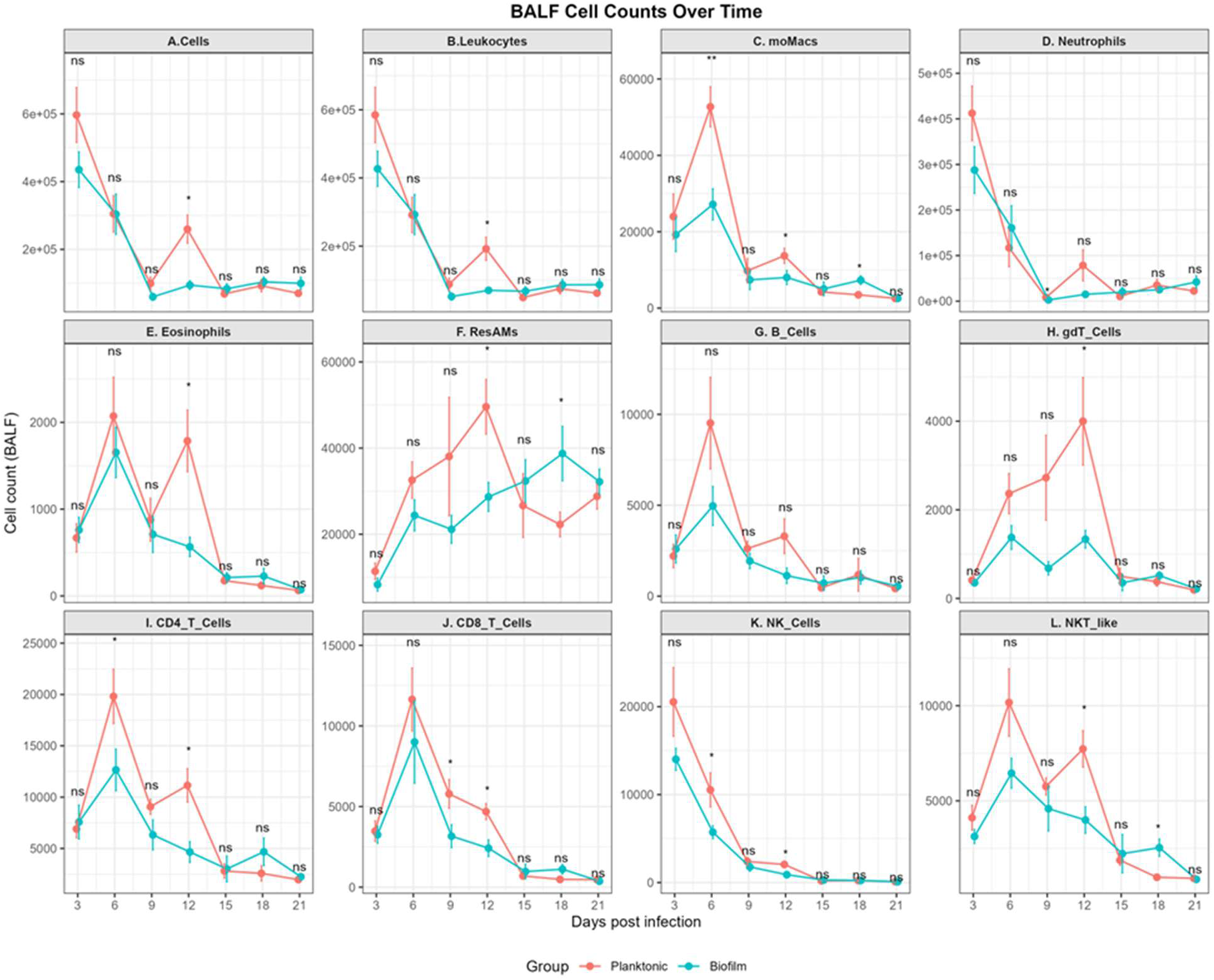
Kinetics of cell counts in bronchoalveolar lavage fluid of *M. pneumoniae* infected mice per day post infection.

**Supplementary Table S1.**
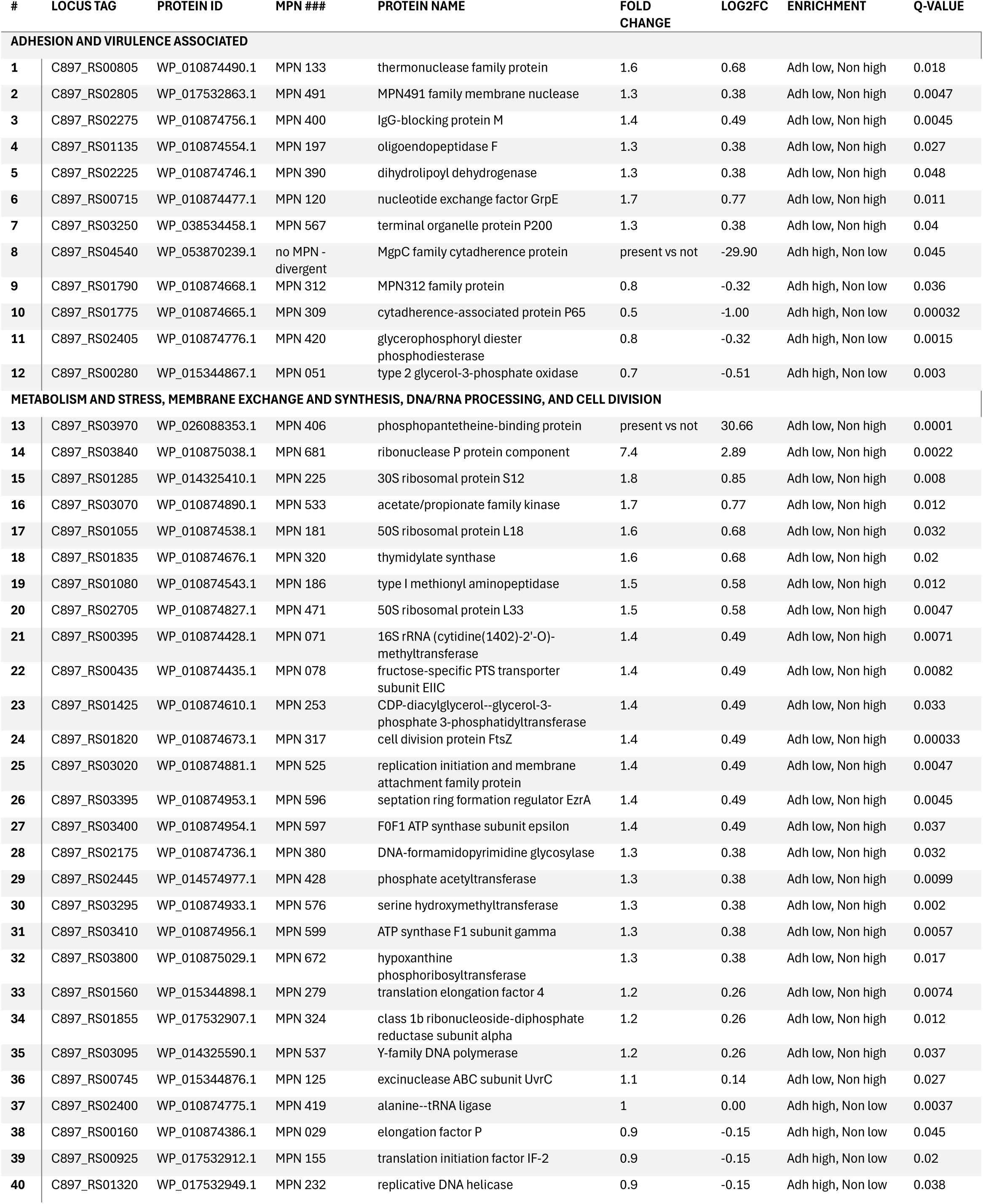
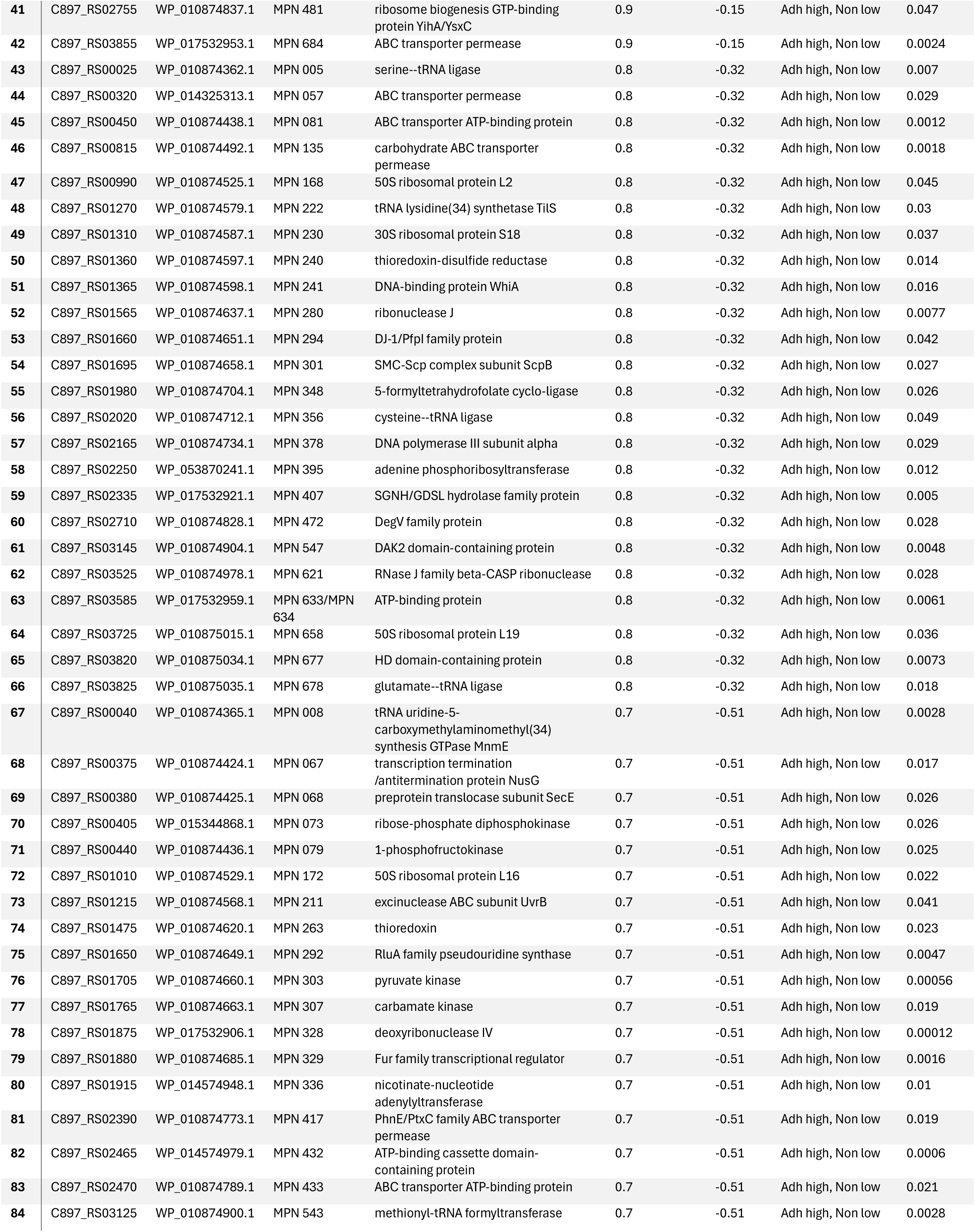
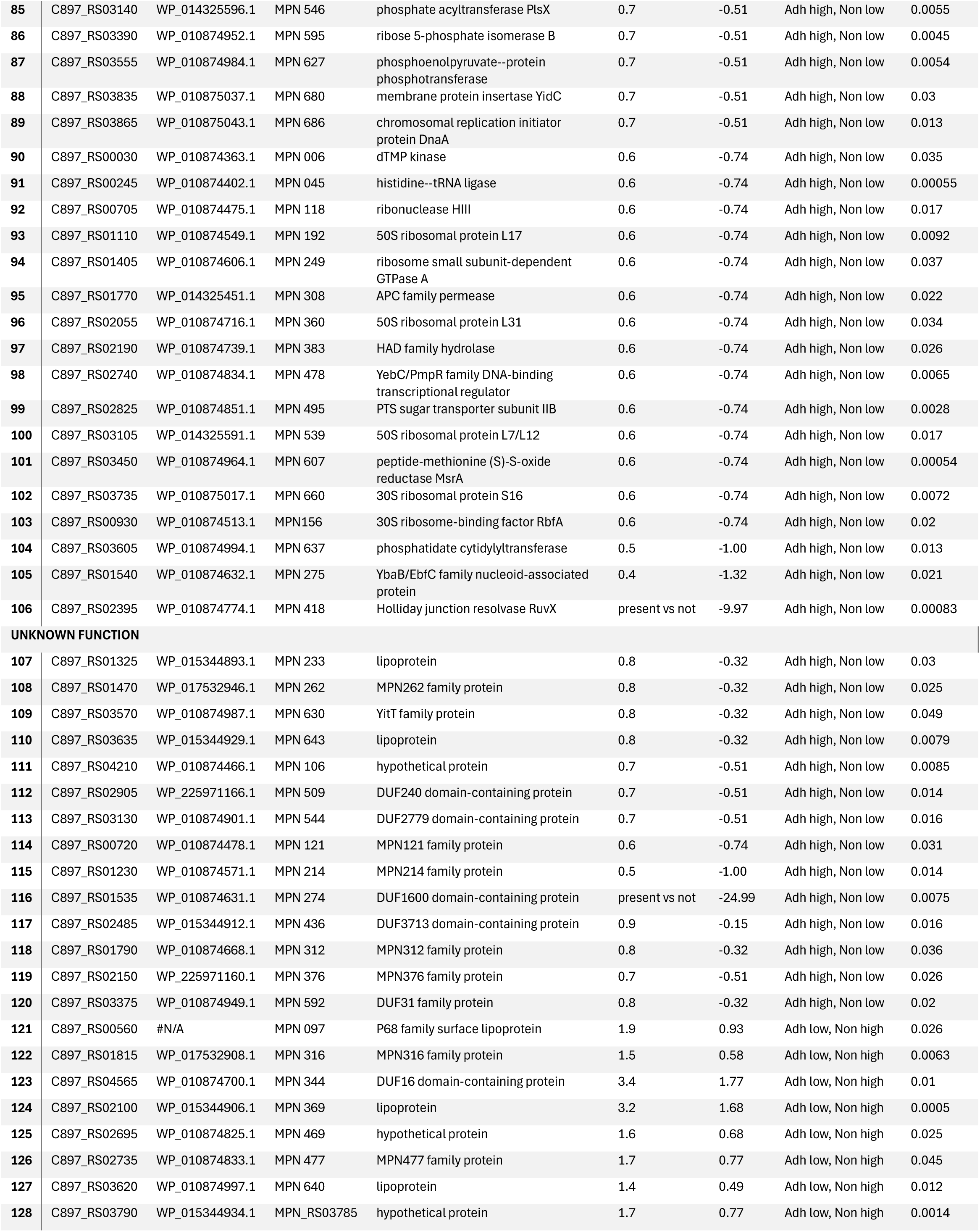
Proteins with statistically significant fraction-specific enrichment.

## Supplementary Proteomics Discussion

### Proteins enriched in the B-Mp Fraction

Several proteins enriched in the B-Mp lack defined functions, including MPN106, MPN121, MPN214, MPN233, MPN262, MPN274, MPN509, MPN544, MPN630, and MPN643. Several proteins involved in adhesion were also upregulated in B-Mp, such as MgpC family cytadherence protein, the MPN213 family protein (MPN312, P24) and the cytadherence-associated protein P65 (MPN309)^51–54^. A DUF31 domain containing protein (MPN592) was also upregulated in B-Mp. Although the function of this specific gene is unknown, DUF31 domain containing proteins have been implicated in the immunoglobulin binding and cleaving MIB-MIP system of some Mycoplasmas, and are believed to possess serine-protease activity^55^.

Multiple proteins involved in membrane exchange were enriched in B-Mp, including ABC transporter permeases (MPN057, MPN081, MPN135, MPN417, MPN432, MPN433, MPN633/634,

MPN684), an APC family permease (MPN308), the membrane protein insertase YidC (MPN680), the preprotein translocase SecE (MPN068), an MPN376 family protein, and a DUF3713 domain-containing protein MPN436. YidC functions as a membrane insertion chaperone that works with the Sec translocon to insert proteins into the membrane^56,57^. SecE also has a defined role in facilitating protein secretion^58^. MPN436 is involved in extracellular protein folding and is crosslinked to MPN376, mutants of which display considerably reduced gliding motility^59,60^.

Membrane synthesis proteins were enriched in B-Mp, including the DegV family protein (MPN372), phosphate acyltransferase PslX (MPN546), and phosphatidate cytidylyltransferase (MPN637, a.k.a CDP-diacylglycerol synthases, CDS). DegV family proteins are known to bind fatty acids, which may imply an upregulation of membrane synthesis^61^. PlsX is a key enzyme in phospholipid and fatty acid formation in *Enterococcus faecalis*, and CDS enzymes are essential for phosphatidylglycerol and cardiolipin synthesis^62,63^.

Multiple components of the phosphoenolpyruvate:sugar phosphotransferase (PEP:PTS) system and glycolysis were enriched in the B-Mp, including phosphoenolpyruvate-protein phosphotransferase (MPN627), the PTS sugar transporter subunit IIB (MPN495), 1-phosphofructokinase FruK (MPN079), pyruvate kinase (MPN303), and dihydoxyacetone kinase DhaK (MPN547) ^64,65^. Nicotinate-nucleotide adenylyltransferase (MPN336), required for NAD+ biosynthesis that feeds into glycolysis, were also enriched in the B-Mp^66,67^. In the biofilm-forming pathogen *Klebisiella pneumoniae*, overexpression of PEP:PTS components enhances biofilm formation by increasing capsular polysaccharide and eDNA production, and it is possible they are serving the same role in *M. pneumoniae* biofilms^68^. MPN303 has also been shown to bind plasminogen, and may contribute to adherence of biofilms on biological surfaces^69^.

Glycerol metabolism and oxidative stress-response proteins were also enriched in B-Mp, including GlpQ (MPN420), GlpO (MPN051), MsrA (MPN607), thioredoxin (MPN240), thioredoxin-disulfide reducatase (MPN263), a Fur family transcriptional regulator (MPN329), a DJ-1/Pfpl family protein (MPN294), the YbaB/EbfC nucleoid-associated protein (MPN275), deoxyribonuclease IV (MPN328) and exinuclease ABC subunit UvrB (MPN211). GlpQ converts glycerophosphodiesters obtained from host membranes into alcohols and glycerol-3-phosphate and is upregulated during nutritional stress in other bacteria^70,71^. GlpO is also upregulated during nutritional stress and utilizes glycerol-3-phosphate to generate DHAP for glycolosis. This procudes H_2_O_2_ as a byproduct^64^. Glycerol metabolism via GlpQ/GlpO has been linked to biofilm formation in *Pseudomonas aeruginosa*, and H_2_O_2_ production has been shown to increase biofilm formation in *Acinetobacter spp.* by inducing exopolysaccharide production^64,72,73^.

MsrA, thioredoxin, and thioredoxin-disulfide reductase mediate oxidative stress defense and redox signaling; MsrA also influences adhesion in a related *Mycoplasma* species^74,75^. Fur regulators mediate metal acquisition (iron, zinc, manganese, nickel) and oxidative stress responses, and zinc metabolism is closely tied to redox homeostasis^76,77^. DJ-1/Pfpl proteins function in stress response, chaperone activity, and proteolysis, while MPN275 binds DNA, alters global transcription, and protects DNA during stress^78,79^. MPN328 and UvrB participate in DNA base excision repair and SOS responses^80,81^.

DNA metabolism, synthesis, and chromosome separation factors were also enriched in B-Mp. These include dTMP kinase TMPK (MPN006), ribose-phosphate diphosphokinase (MPN073), ribose-5- phosphate isomerase B (MPN595), adenine phosphoribosyltransferase (MPN395), the replication initiator DnaA (MPN686), the replicative helicase DnaB (MPN232), DNA polymerase III subunit alpha (MPN378), Holliday junction resolvase RuvX (MPN418), the SMC-Scp complex subunit ScpB (MPN301), and the DNA-binding protein WhiA (MPN241). TMPK, MPN073, MPN595, and MPN395 all contribute to nucleotide metabolism by participating in pyrimidine synthesis (TMPK), functioning in the pentose phosphate pathway (MPN073, MPN595), and adenine salvaging for nucleotide synthesis (MPN395)^82–85^. DnaA, DnaB and MPN378 mediate DNA replication^86–88^. RuvX, ScpB and WhiA contribute to DNA separation, with ScpB and WhiA being required for chromosome segregation^89,90^. RuvX participates in homologous recombination, repair of collapsed/stalled forks, and chromosome segregation. It is often upregulated during DNA damage, potentially reflecting oxidative stress or preparation for division^91^. In the closely related *Mycoplasma genitalium*, the RuvABC system drives antigenic variation of the MgpB and MgpC adhensins through recombination with archived gene copies. Notably, *M. pneumoniae* lacks a functional RuvC, so RuvX may be compensating for this role. In other bacteria, RuvC has been shown to process extracellular DNA (eDNA) in biofilms, a critical component of the extracellular polymeric substance (EPS) matrix in bacterial biofilms^92–94^. While evidence of eDNA involvement in *M. pneumoniae* biofilms is limited, eDNA is an essential component for *Mycoplasma hyopneumoniae* biofilm formation, and it is possible that this is the case for *M. pneumoniae* as well^94^.

Proteins involved in RNA metabolism were also enriched in B-Mp. These include multiple aminoacyl-tRNA ligases (serine MPN005, histidine MPN045, cysteine MPN356, alanine MPN419, and glutamate MPN678,) as well as 30S ribosome-binding factor RbfA (MPN156), 30S ribosomal proteins S16 (MPN660) and S18 (MPN230), 50S ribosomal proteins L2 (MPN168), L7/L12 (MPN539), L16 (MPN172), L17 (MPN192), L19 (MPN658), and L31 (MPN360). Additional upregulated factors include translation intitation factor IF-2 (MPN155), ribosome-associated GTPases (MPN249, MPN481), methionyl-tRNA formyltransferase (MPN543), elongation factor P (MPN029), tRNA lysidine synthetase TilS (MPN222), transcription termination/antitermination protein NusG (MPN067), GTPase MnmE (MPN008), RluA family pseudouridine synthase (MPN292), ribonuclease J (MPN280), RNase J family β-CASP ribonuclease (MPN621), and ribonuclease HIII (MPN118). MPN292 catalyzes pseudouridylation of tRNAs and rRNAs, stabilizing RNA duplexes^95^. MPN280 and MPN621 are RNAse J family enzymes with exo- and endonuclease activity involved in RNA processing, and MPN 118 degrades RNA in RNA-DNA hybrids^96,97^.

Proteins with diverse functions were enriched in the B-Mp, including an HD-domain-containing protein (MPN677), a HAD family hydrolase (MPN383), and a YebC/PmpR family DNA-binding transcriptional regulator (MPN378). HD-domain proteins are metalloproteins that have been shown to catalyze reactions ranging from hydrolysis to oxygenation, while YebC regulators influence virulence associated processes such as surface antigen expression in *Borrelia burgdorferi*, T3SS expression in *Edwardsiella piscicida*, and surface polysaccharide expression in *E. coli*^98–101^. HAD-family proteins are a very large family of proteins playing a multitude of roles in organisms of various orders. Specific members have been strongly implicated in the virulence of various bacteria (10.1016/j.abb.2024.109888).

Finally, enriched in B-Mp were proteins strongly implicated in biofilm formation, including an SGNNH/GDSL hydrolase family protein (MPN407), carbamate kinase (MPN307), and 5-formyltetrahydrofolate cyclo-ligase (MPN348). SGNH hydrolases function as carbohydrate-modifying enzymes, and homologs such as AlgX in *Pseudomonas aeruginosa* are directly required for biofilm formation^102,103^. MPN307 participates in arginine metabolism which has been linked to biofilm formation in other bacteria, although this pathway is believed to be non-functional in M. pneumoniae^104,105^. MPN348 is an ATP-dependent folate salvage enzyme that is often upregulated in bacterial biofilms^106,107^.

### Proteins enriched in the P-Mp fraction

Much like what was observed for the B-Mp fraction, many proteins enriched in the P-Mp fraction (MPN097, MPN316, MPN344, MPN369, MPN469, MPN477, MPN484, MPN640, and MPN_RS03785) lack known functions. These factors should be the subject of further research, as their enrichment in the more virulent P-Mp fraction implies they may serve as potential virulence factors. Classically recognized *M. pneumoniae* virulence factors were also enriched in the P-Mp. These include the thermonuclease family protein (MPN133), the membrane nuclease (MPN491), the IgG-blocking protein M IbpM (MPN400), oligoendopeptidase F (MPN197), dihydrolipoyl dehydrogenase (MPN390, DLD, DLDH, lipoamide dehydrogenase, *lpd*, E3*, pdhD*), nucleotide exchange factor GrpE (MPN120), and P200 (MPN567).

MPN133 is a cytotoxic nuclease that can be internalized into cells and cause cell death and pathology and MPN491 is a secreted nuclease that mediates immune evasion by digesting neutrophil extracellular traps^108,109^. IbpM mediates antibody escape by sequestering IgM, IgG and IgA antibodies, and a mutant in this gene has displayed reduced cytotoxic capabilities^110^. MPN197 is a surface accessible peptidase that may be involved in *M. pneumoniae* surface protein presentation and has been found to serve as a negative regulator of biofilms^111,112^. A homolog of this protein in a related Mycoplasma species was found to cleave bradykinin, substance P, neuropeptide Y, and neurokinin A, potentially implicating it as a virulence factor as well^113^. MPN390 is a critical enzyme involved in the pyruvate dehydrogenase complex and has been demonstrated to be a key virulence determinant for multiple organisms, including the closely related *Mycoplasma gallisepticum*. It has also been identified as a plasminogen-binding protein and is thus also implicated in adherence^114–117^. GrpE is a co-chaperone with DnaK and has been found to be located on the surface of other Mycoplasma species and contribute to adhesion by interacting with plasminogen and other extracellular matrix components^118^. Further demonstrating the increased ability of P-Mp to adhere to host components, is the upregulation of the adherence associated protein P200 (MPN567) which is required for attachment to differentiated bronchial epithelium and for the gliding motility of the organism^119^. The enriched P200 in P-Mp relative to B-Mp is consistent with other literature on *M. pneumoniae* biofilms indicating that P200 is downregulated over time as biofilms mature^44^.

Moving away from the virulence axis and more on the metabolic phenotypes, proteins associated with ATP production were enriched in P-Mp, identifying this fraction as one that is metabolically active.

These proteins include F0F1 ATP synthase subunit epsilon (MPN597) and ATP synthase F1 subunit gamma (MPN599)^120^. Also enriched in P-Mp were proteins associated with cell division, including replication initiation and membrane attachment family protein (MPN525), FtsZ (MPN317), and EzrA (MPN596). MPN525 is highly homologous to DnaD proteins that are involved in the primosomal step of DNA replication, while FtsZ and EzrA are involved in Z-ring formation of dividing cells^121,122^. Proteins involved in DNA metabolism and repair were also upregulated, including class 1b ribonucleoside- diphosphate reducates subunit alpha (MPN324, RNR), hypoxanthine phosphotibosyltransferase (MPN672, HPRT), exinuclease ABC subunit UvrC (MPN125), DNA-formamidopyrimidine glycosylase (MPN380) and Y-family DNA polymerase (MPN537). RNRs are responsible for converting ribonucleotides to deoxyribonucleotides and HPRT is an enzyme involved in purine scavenging^123,124^. UvrC is involved in excision of damaged sites during nucleotide excision repair and MPN380 recognizes oxidized bases and is involved in base excision repair^80,125^. Y-family polymerases are involved in translesion synthesis in damaged DNA^126^.

Also enriched in P-Mp were proteins involved in metabolic changes implicated in switching between different phenotypes. These include thymidylate synthase (MPN320, TS), serine hydroxymethylastransferase (MPN576, SHMT), phosphate acetyl transferase (MPN428, PTA) and acetate/proprionate family kinase (MPN533, AckA). TS and SHMT are heavily involved in 1-carbon metabolism and DNA synthesis^127^. TS converts dUMP into dTMP, and intracellular dTMP levels have been found to modulate cell phenotypes in other bacteria where low levels lead to small-cell variants that have improved intracellular survival^128^. SHMT is also implicated in biofilm formation, as a *P. aeruginosa* SHMT mutant increased its biofilm formation by 45-fold and resulted in a small-colony variant of the pathogen^129^. The observed combination of PTA and AckA was also notable because the PTA-AckA pathway operates as a carbon overflow pathway that is associated with fast growth states typically associated with planktonic fractions. Furthermore, when an intermediate within the pathway, acetyl phosphate, builds up, it is reported to induce the transition from planktonic states toward biofilm formation^130^. Also upregulated in P-Mp are the fructose-specific PTS transporter subunit EIIC FruA (MPN078), phosphopantetheine-binding protein (MPN406, acyl carrier protein) and CDP-diacylglycerol-glycerol-3phosphate-3-phosphatidyltransferase (MPN253). Fructose transported in by FruA can be used to undergo multiple metabolic pathways^64^. MPN253 is reported to help in synthesizing acidic phospholipids^131^.

Noteworthy was the significant upregulation of the *phosphopantetheine binding protein* (AcpP, MPN406) in P-Mp, which was almost entirely absent in the B-Mp. MPN406 is an acyl carrier protein that is a component of fatty acid synthesis (FAS), non-ribosomal polypeptide synthesis (NRPS), and polyketide synthesis(PKS)^132^. Acyl carrier proteins are implicated in bacterial virulence through 4’- phosphopantetheinyl transferases (PPTases), which are enzymes that post-translationally attach a phosphopantetheine moiety (Ppant) from CoA onto carrier proteins. This attachment activates these carrier proteins for biosynthesis of virulence-related molecules in other bacterial pathogens such as *Mycobacteria* spp. *Salmonella enterica*, pathogenic *E. coli*, *Burkholderia*, *Acinetobacter*, *Pseudomonas aeruginosa,* and even some fungi^133–146^. *M. pneumoniae* lacks the ability to synthesize fatty acids *de novo,* and instead extracts them from the host cells through the actions the lipid scavenger protein P116 (MPN_213)^147^. However, *M .pneumoniae* still possesses the acyl carrier protein synthase (AcpS |MPN_298) which is the 4’-phosphopantetheinyl transferase (PPTase), and the acyl carrier protein AcpP (MPN406) which provides phosphopantetheine as a cofactor for delivery acyl groups to phospho- or glycolipids, which these genes may be involved in the synthesis of^133,146^. Both genes are suggested to be essential, and data are limited on whether they play a role in the virulence of *M. pneumoniae*. *M. pneumoniae* also lacks the genes required for the synthesis of traditional siderophores, and it is not currently known whether the production of the CARDS toxin or other *M. pneumoniae* virulence factors relies on AcpP activity. However, many Mycoplasma proteins are known to moonlight as virulence factors and given that the AcpS/AcpP axis is associated with *in vivo* virulence of other bacterial pathogens, it is tempting to speculate that AcpP may contribute to, either directly or indirectly, to the heightened virulence of P-Mp.

Also enriched in P-Mp were multiple proteins involved in translation, ribosome function and tRNA processing. These include 30S ribosomal protein S12 (MPN225), 50S ribosomal protein L18 (MPN181), 50S ribosomal protein L33 (MPN471), translation elongation factor 4 (MPN279), type I methionyl aminopeptidase (MPN186), 16S rRNA (cytidine(1402)-2’-O)-methyltransferase (MPN071), and ribonuclease P protein component (MPN681).

